# Functional determinants and evolutionary consequences of pleiotropy in complex and Mendelian traits

**DOI:** 10.1101/2024.10.28.620594

**Authors:** Yury A. Barbitoff, Polina M. Bogaichuk, Nadezhda S. Pavlova, Alexander V. Predeus

## Abstract

Pleiotropy, a phenomenon of multiple phenotypic effects of the same genetic alteration, is one of the most important features of genotype-to-phenotype networks. Over the last century, biologists have actively debated the prevalence, mechanisms, and consequences of pleiotropy. In this work, we employed data on genotype-to-phenotype associations from the Human Phenotype Ontology and Mouse Genome Database, as well as genome-wide associations from the UK Biobank cohort to investigate the similarities and dissimilarities in the patterns of pleiotropy between species and different trait types (i.e., Mendelian traits and complex traits). We found that the pleiotropic effects of genes correlate well between species, but have a much weaker correlation when different data sources are compared for the same species. In all cases, however, highly pleiotropic genes possessed a common set of features, such as the broad expression across tissues, involvement in many biological processes, or a high number of protein-protein interactions of the respective gene products. Furthermore, we observed a universal tendency of highly pleiotropic genes to be under greater negative selection pressure combined with a significant enrichment of recent positive selection signals at pleiotropic loci. Taken together, our results pinpoint a common mechanism underlying pleiotropic effects in different trait domains, and suggest that high degree of pleiotropy plays a role in adaptation, despite imposing additional constraint on genetic variation.

## Introduction

Reconstruction of the complex network of genotype-to-phenotype relationships is a pivotal task of modern genetics. Over the recent years, development and application of high-throughput methods has facilitated the construction of large-scale resources that aggregate data on genotype-to-phenotype associations, such as the Mouse Genome Database (MGD) (Blake et al., 2003) or the National Human Genome Research Institute (NHGRI) GWAS Catalog (Buniello et al., 2018). Construction of the phenotype ontologies, such as the Human Phenotype Ontology (Robinson et al., 2008) and the Mammalian Phenotype Ontology (MP) (Smith et al., 2004), has further enabled accurate integration and comparison of genotype-to-phenotype relationships, both within and between species.

Pleiotropy is a phenomenon in which a single mutation affects multiple independent traits in the phenotype (Wagner and Zhang, 2011), and is thus one of the major forces that shapes the genotype-to-phenotype networks. Since the advent of genomics, multiple studies have demonstrated the abundance of pleiotropic effects in different species, and different theoretical models of pleiotropy have been proposed. The modular pleiotropy model predicts that genes with similar functions should affect groups (or “modules”) of related traits, while the universal pleiotropy model (also known as the omnigenic hypothesis) states that all genes can be expected to impact all traits (reviewed in Tyler et al., 2016). Recent studies favor the modular pleiotropy model, with two key pieces of supporting evidence: (i) the observed degree of pleiotropy for most mutations is modest, and (ii) the networks of gene-trait connections have a high degree of modularity. For example, both of these features of genotype-to-phenotype networks have been demonstrated in a comparative study of yeast, nematodes, and mice (Wang et al., 2010).

Genome-wide association studies (GWAS) provide a rich set of information for studying pleiotropy in the human genome, and multiple such studies have been performed (e.g., Pickrell et al., 2016; Chesmore et al., 2018). Emergence of large-scale deeply phenotyped cohorts, such as the UK Biobank (UKB) (Bycroft et al., 2018), has opened new prospects for pleiotropy research. Several efforts have been undertaken to study pleiotropy in UKB (Watanabe et al., 2019; Jordan et al., 2019), including a study performed by our group (Shikov et al., 2020). These studies have confirmed that pleiotropic effects are typical for a large proportion of genes, although the estimated proportion of pleiotropic loci varied widely between studies.

Estimation of the degree of pleiotropy from GWAS data is complicated by various factors, including (i) linkage disequilibrium (LD) between multiple genetic variants at a locus, leading to spurious pleiotropy (Chebib et al., 2021); (ii) genetic and environmental correlations between traits that inflate the estimated level of pleiotropy; and (iii) the burden of multiple testing of association between millions of variants and thousands of traits (reviewed in Hackinger and Zeggini, 2017). In line with these considerations, the estimated proportion of loci with pleiotropic effects in the UKB data was heavily dependent on the approach to defining orthogonal (independent) phenotypes. For example, more than half of the human genome was reported to bear signals of pleiotropy by Watanabe et al. (Watanabe et al.,2 019) when using trait domain as a measure of trait independence; however, our analysis based on hierarchical clustering of traits by their phenotypic correlation yielded a much more conservative estimate of 5.4% of the genome (Shikov et al., 2020).

A particularly intriguing question that arises when analyzing pleiotropic effects of genetic variation is the evolutionary consequences of pleiotropy (reviewed in Zhang, 2023). In 2000, Orr found that pleiotropy should naturally decrease the rate of adaptation within the Fisher’s geometric model framework (Orr, 2000). This model, however, assumes constant total effects of mutations, and contradicts empirical observations (e.g., Wagner et al., 2008). Furthermore, Wang et al. have shown that pleiotropy could, to the contrary, increase the rate of adaptation if the effect size of a mutation is set to be dependent on the degree of pleiotropy (Wang et al., 2010). Further support for this model comes from studies that report an enrichment of pleiotropic loci in humans with signals of recent positive selection (e.g. Sakaue et al., 2021). The adaptive effect, however, is expected to be restricted to loci with moderate degree of pleiotropy, as evidenced by studies in plants (Frachon et al., 2017).

In our earlier work, we have observed an unexpected enrichment of common variants at pleiotropic loci, an observation that challenges the concept of the evolutionary cost of pleiotropy and could possibly indicate an adaptive role of pleiotropic genetic variants. A later study by Novo et al. suggested that pleiotropic loci are subject to strong background selection; however, the significance of the trend varied for different source data were used (Novo et al., 2021). Furthermore, the *B* statistic used by Novo et al. can deviate from expectation due to different selection types, and may also be confounded by recombination rate. Taken together, results of previous studies do not allow a definitive conclusion regarding the evolutionary consequences of pleiotropy in the mammalian genome.

To address this issue, as well as to provide a deeper insight into the mechanistic basis of pleiotropy in mammals, we have jointly analyzed pleiotropic effects of human and mouse genes using both Mendelian trait data (provided by the Human Phenotype Ontology), Mouse Genome Database), and biobank-scale phenome-wide association studies (UK Biobank) to analyze functional properties of pleiotropic genes in various species and trait domains.

## Results

### Evaluating the concordance of pleiotropic effects between species and trait domains

To compare the patterns of pleiotropy between species and trait domains, we first set off to construct a dataset based on publicly available information on genotype-to-phenotype relationships in humans and mice. To this end, we collected and merged data from three main sources: the Human Phenotype Ontology (HPO), the Mouse Genome Database (MGD), and the pan-UK Biobank (pan-UKB) study. The former two sources contain human (HPO) and murine (MGD) gene annotations with human or mammalian phenotype ontologies, respectively, while the latter dataset represents a compendium of genome-wide association results for as many as 1263 human complex traits (see Methods and Supplementary Note for details on data pre-processing). In total, the dataset contained 150,053 gene-to-trait relationships, with 53,406, 82,715, and 13,932 coming from each of the aforementioned data sources. A total of 4,999, 13,301, and 6,649 genes were associated with at least one trait in HPO, MGD, and pan-UKB data, respectively. Of note, only 1,236 genes had at least one gene-trait association across all three datasets, and trait associations from both trait domains (from pan-UKB and either HPO or MGD) were available for 3,188 genes. Thus, more than half of the gene-trait connections were unique to each trait domain (52.0% for complex traits data, and 77.1% - for Mendelian trait data).

Before delving into the comparative analysis of the obtained data, we first examined the distribution of the degree of pleiotropy observed in each dataset. While all distributions had a characteristic left skew (Figure 1a), the overall shape of the distribution varied. The distribution was much more asymmetric in case of the genome-wide association data (Figure 1a, right plot), with a median of 1 trait clusters associated with a gene. On the other hand, in both HPO- and MGD-based data, the median number of phenotype terms associated with each gene was much greater (11 and 5, respectively), corresponding to a lesser skew of the distribution (Figure 1a, left and center plots). Only 287 out of 4999 human genes (5.7%) were associated with exactly 1 upper-level HPO term, i.e. were non-pleiotropic (for comparison, 56.4% of genes were non-pleiotropic in GWAS). In MGD, non-pleiotropic genes comprised a slightly greater fraction of genes (1409 out of 13,301, 10.6%). In line with the differences in the distribution shape, the genotype-to-phenotype networks differed in the estimated degree of modularity (0.118, 0.205, and 0.516 for HPO, MGD, and pan-UKB data, respectively), with complex trait-based network having by far the greatest level of modularity.

**Figure 1.**
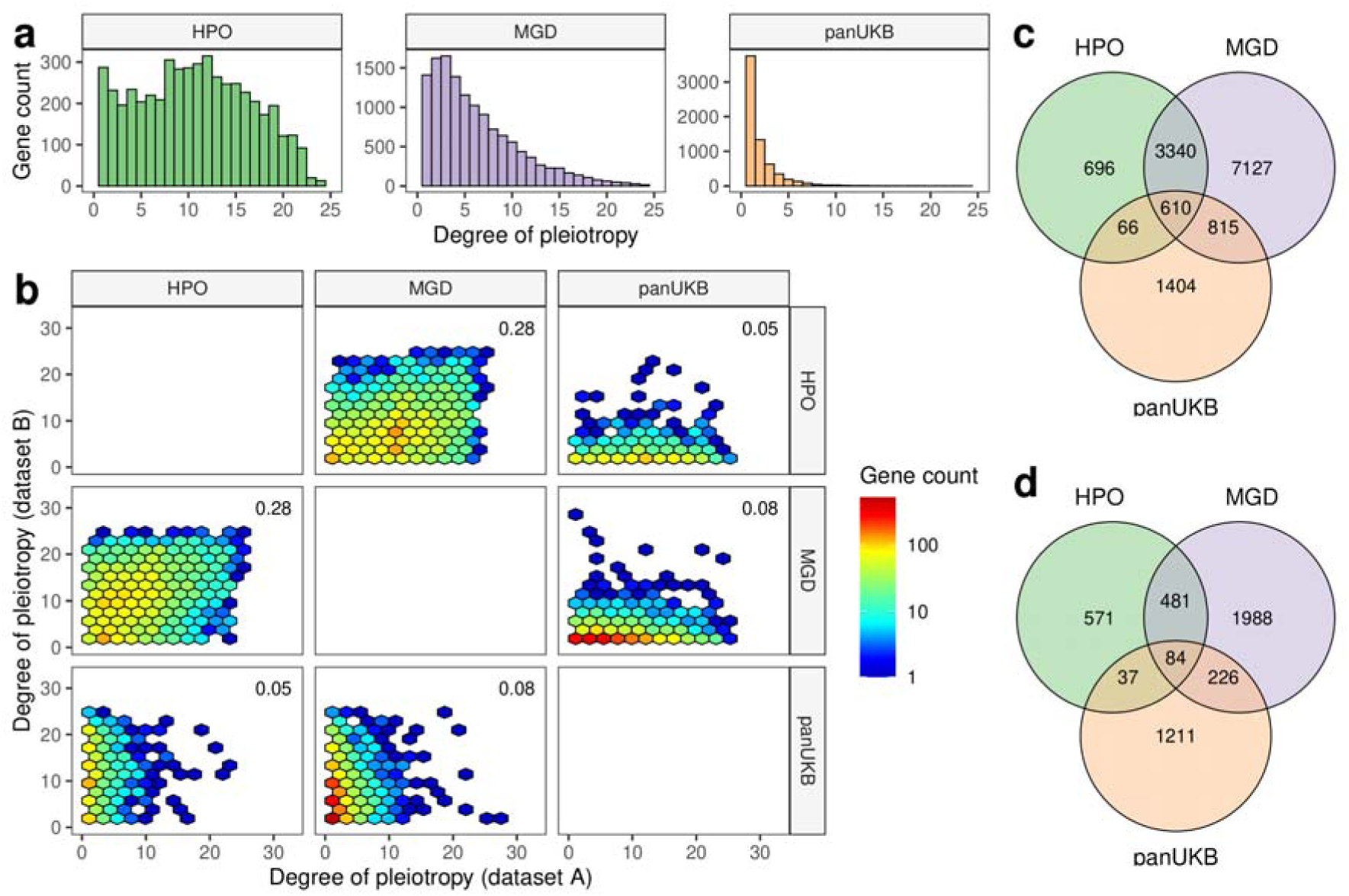
Gene-level degrees of pleiotropy correlate between species and trait domains. (a) Distribution of the degree of pleiotropy of human and mouse genes according to different sources of data (from left to right - Human Phenotype Ontology, Mouse Genome Database, pan-UK Biobank GWAS data. (b) Hexagonal scatter plots showing the correspondence between the degree of pleiotropy of genes between different data sources. (c, d) Venn diagrams showing the overlaps between sets of all pleiotropic genes (c) or highly pleiotropic genes (d) between the three datasets (see text for details on cutoff selection).

Having characterized the overall structure of the data, we next went on to compare the patterns of pleiotropy between the datasets. To this end, we used orthologous gene mapping between human and mouse to create a single merged dataset. Comparison of the degree of pleiotropy for single-copy orthologous gene pairs in mouse and human revealed a modest significant correlation (Spearman’s p = 0.28, Figure 1b). At the same time, the degree of pleiotropy estimated using complex trait data and Mendelian trait data showed much lower correlation. For instance, there was no significant correlation between the gene pleiotropy in HPO and pan-UKB (Figure 1b). Remarkably, the degree of pleiotropy estimated for murine genes using MGD data significantly correlated with GWAS-based estimates for human orthologs (Spearman’s correlation = 0.08, p < 0.001). The greater significance of the estimates for MGD data compared to HPO may be explained by the higher number of genes with non-zero degree of pleiotropy in mice.

Given the relatively low correlation between the degree of pleiotropy estimates, we next questioned if a different result could be obtained when testing the overlap between sets of pleiotropic and non-pleiotropic genes without taking into account the numeric estimate of the degree of pleiotropy. To this end, we divided each dataset into three categories: (i) non-pleiotropic genes (associated with 1 upper-level ontology term or 1 complex trait cluster); (ii) highly pleiotropic genes (identified as top quartile of genes by degree of pleiotropy; the trait/term count cutoffs are 15 (HPO), 9 (MGD), and 2 (pan-UKB data); and (iii) modestly pleiotropic genes (this group includes all of the remaining pleiotropic genes). The proportions of genes in different groups can be seen on Extended Data Figure 1.

We next compared the sets of pleiotropic genes between the dataset. When moderately pleiotropic and highly pleiotropic genes were combined, a total of 610 genes overlapped between all datasets (Figure 1c). When only highly pleiotropic genes were considered, 84 genes emerged as overlapping between all data types (Figure 1d). Importantly, as many as 3340 genes were pleiotropic according to both HPO and MGD data, but were not pleiotropic in GWAS, and a total of 11,163 genes were uniquely pleiotropic according to either HPO or MGD data. This amounts to 88.2% of all genes that were pleiotropic in either human or mouse based on Mendelian trait data. At the same time, only 48.5% (1404) of genes pleiotropic in pan-UKB data were not pleiotropic in the Mendelian trait domain, with 1307 of these genes having no phenotype associations in either HPO or MGD. Of note, the percentages of shared highly pleiotropic genes was even lower than those observed for all pleiotropic genes (only 10.5% of genes highly pleiotropic in HPO or MGD were pleiotropic in GWAS).

Taken together, these numbers suggest that, despite low correlation between the degrees of pleiotropy between complex and Mendelian traits, there is a significant overlap between the sets of pleiotropic genes between trait domains. It is also important to note that human orthologs of as many as 815 genes with pleiotropic effects in MGD, but not HPO data, were pleiotropic according to pan-UKB GWAS data. These findings may suggest that many of the genes that demonstrate large effects on the phenotype in mice drive complex rather than Mendelian traits in humans.

### Pleiotropic genes are characterized by common functional properties

Having split the dataset into different groups of genes depending on their estimated degree of pleiotropy, we next sought to identify shared features of pleiotropic genes between species and trait domains. To do so, we have annotated our dataset with various gene-level properties, such as the summary statistics of gene expression profile (human gene data from GTEx were used), numbers of protein-protein interactions (according to the STRING database), and measures of haploinsufficiency and triplosensitivity (Collins et al., 2022).

Analysis of the expression pattern of genes in different pleiotropy group showed that more pleiotropic genes tend to have broader expression profile (as evidenced by the number of tissues with expression at the level of at least 5 TPM (Wilcoxon-Mann-Whitney test p-value < 0.001 when comparing non-pleiotropic and highly pleiotropic genes in either of the data sources; Figure 2a) and greater mean/median expression across tissues (Wilcoxon-Mann-Whitney test p-value < 0.001, Extended Data Figure 2)). Importantly, the number of tissues with detectable expression was much greater for highly pleiotropic genes in the Mendelian trait data (30 tissues for HPO, and 27 - for MGD) compared to those identified from GWAS data (9). Besides the breadth of expression across tissues, we also observed a significant increase in the maximum expression level of genes with greater degrees of pleiotropy (p < 0.001 in all cases, Extended Data Figure 2). This difference was the most pronounced for genes at pleiotropic GWAS loci (median of highest expression values - 22.2, 28.4, 36.3 TPM for non-pleiotropic, moderately pleiotropic, and highly pleiotropic genes). Similarly to the results of the breadth of expression profile, pleiotropic Mendelian trait genes had higher expression level compared to genes at pleiotropic GWAS loci (55.3 TPM for HPO data, and 56.8 - for MGD vs. 36.3 for GWAS).

**Figure 2.**
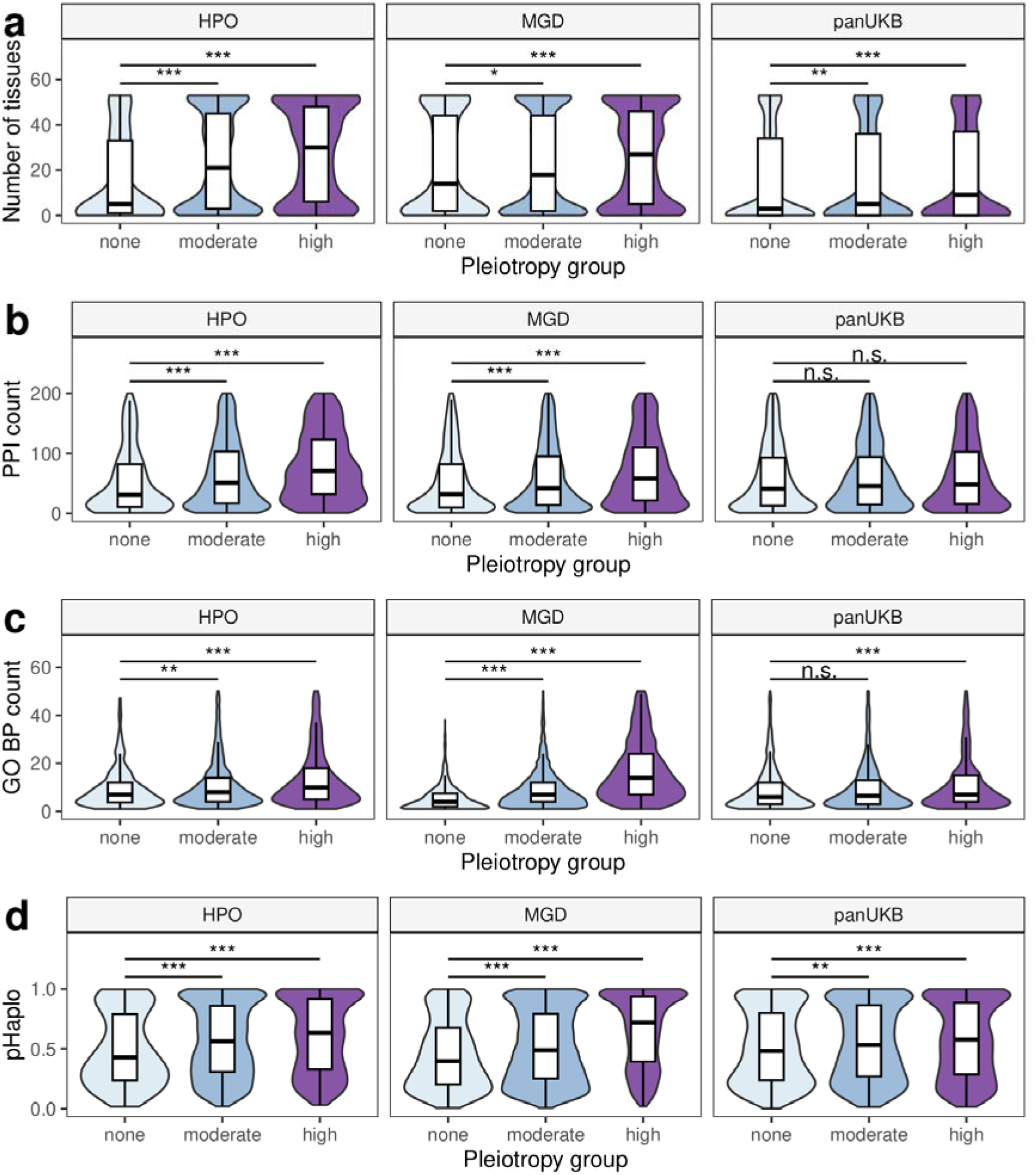
Pleiotropic effects correlate with functional significance of genes in both trait domains. Shown are violin and box plots for (a) number of tissues expressing a gene at the level of at least 5 TPM according to the Genotype Tissue Expression (GTEx) v8 data; (b) number of protein-protein interactions according to the STRING database; (c) number of biological process (BP) terms from Gene Ontology (GO); and (d) pHaplo scores from Collins et al., 2022. ** - p < 0.01, *** - p < 0.001, n.s. - no significant differences in Wilcoxon-Mann-Whitney test with Benjamini-Hochberg FDR adjustment.

Besides broader expression, we also demonstrated that the genes with greater degree of pleiotropy encode proteins with a larger number of protein-protein interactions. This tendency, however, was more apparent in the Mendelian trait domain (median number of interactions for highly pleiotropic proteins = 127 and 101 compared to 42.5 and 45.0 for non-pleiotropic human and murine genes, respectively, p < 0.001) (Figure 2b). For GWAS-based data, no statistically significant difference in the number of PPIs was detected (however, the difference could be seen with a different approach to GWAS data preprocessing, see Supplementary Note, section 2). Again, the number of interactions for highly pleiotropic GWAS genes was much smaller compared to highly pleiotropic Mendelian trait genes, in line with our observations made during expression pattern analysis.

As a next step, we then tested whether the highly pleiotropic genes have broader sets of functions compared to non-pleiotropic ones. To this end, we have annotated each gene with the number of corresponding Gene Ontology (GO) terms, and compared these numbers between gene groups. Indeed, highly pleiotropic genes had higher numbers of associated terms in all three GO branches (biological process (BP), cellular component (CC), and molecular function (MF)) (Figure 2c, Extended Data Figure 3). Thus, a median highly pleiotropic Mendelian trait gene was involved in 10 and 16 biological processes (for human and mouse, respectively, according to HPO and MGD data), compared to only 6 and 4 - for non-pleiotropic ones. In the complex trait domain, the difference was much smaller (8 vs. 6), but still highly significant (p << 0.001).

Given the above observations, we also tested whether the highly pleiotropic genes have higher degree of dosage sensitivity. To perform such a test, we employed the pHaplo and pTriplo metrics calculated in a recent study of copy number variation in the human genome (Collins et al., 2022). Indeed, we observed that more pleiotropic genes have higher pHaplo and pTriplo values (Figure 2d, Extended Data Figure 4). The difference in pHaplo and pTriplo was also weaker for complex trait data (highly pleiotropic vs. non-pleiotropic, pHaplo: 0.58 vs 0.48 in pan-UKB data, 0.62 vs 0.39 in HPO, 0.72 vs 0.40 in MGD; pTriplo: 0.49 vs 0.43 in pan-ULB; 0.54 vs 0.40 in HPO, 0.60 vs 0.38 in MGD).

Besides the analysis of various measurable properties of the genes, we also used gene set enrichment analysis to get insights into the function of the genes in different pleiotropy groups. Due to the inherent uncertainty of causal gene identification from GWAS data, we primarily focused this analysis on genes identified using Mendelian trait data (HPO and MGD). For highly pleiotropic genes in both humans and mice, analysis of canonical pathways and hallmark gene sets demonstrated enrichment for pathways related to cell proliferation and DNA repair (e.g., KEGG’s collection of pathways in cancer, UV response and TGF□ signaling hallmarks) (Extended Data Figures 5-6). Similar results were obtained when analyzing GO term enrichment, with development, morphogenesis, and cell proliferation among the most enriched BP terms (Extended Data Figure 7). In line with these observations, molecular function (MF) term analysis identified terms related to chromatin and DNA binding, transcription factor binding, and protein complex formation to be the most overrepresented (Extended Data Figure 8). Again, these patterns were shared for highly pleiotropic human and murine genes, and both of these sets were highly enriched for cellular component (CC) terms associated with chromatin (Extended Data Figure 9).

Of note, we also found that highly pleiotropic GWAS genes tend to show enrichment for highly similar gene sets. Thus, 125 enriched MSigDB canonical pathways were shared between highly pleiotropic genes from all three data sources, and only a minor fraction of pathways (54, or 24.2%) were uniquely enriched among highly pleiotropic complex trait genes (Figure 3a). The degree of overlap was even more significant for enriched GO terms, with 1094 BP terms shared between all three data sources, and only a handful of terms (28, or 2.0%) were uniquely enriched among pleiotropic GWAS genes. In other GO branches, the proportion of terms specific to pan-UKB was comparably small (12.1% for CC, and 6.1 - for MF, corresponding to 9 terms in each case) (Extended Data Figure 11). These results show that the overlap in the biological processes and molecular functions of highly pleiotropic genes is greater compared to the overlap of the gene sets themselves (Figure 1d).

**Figure 3.**
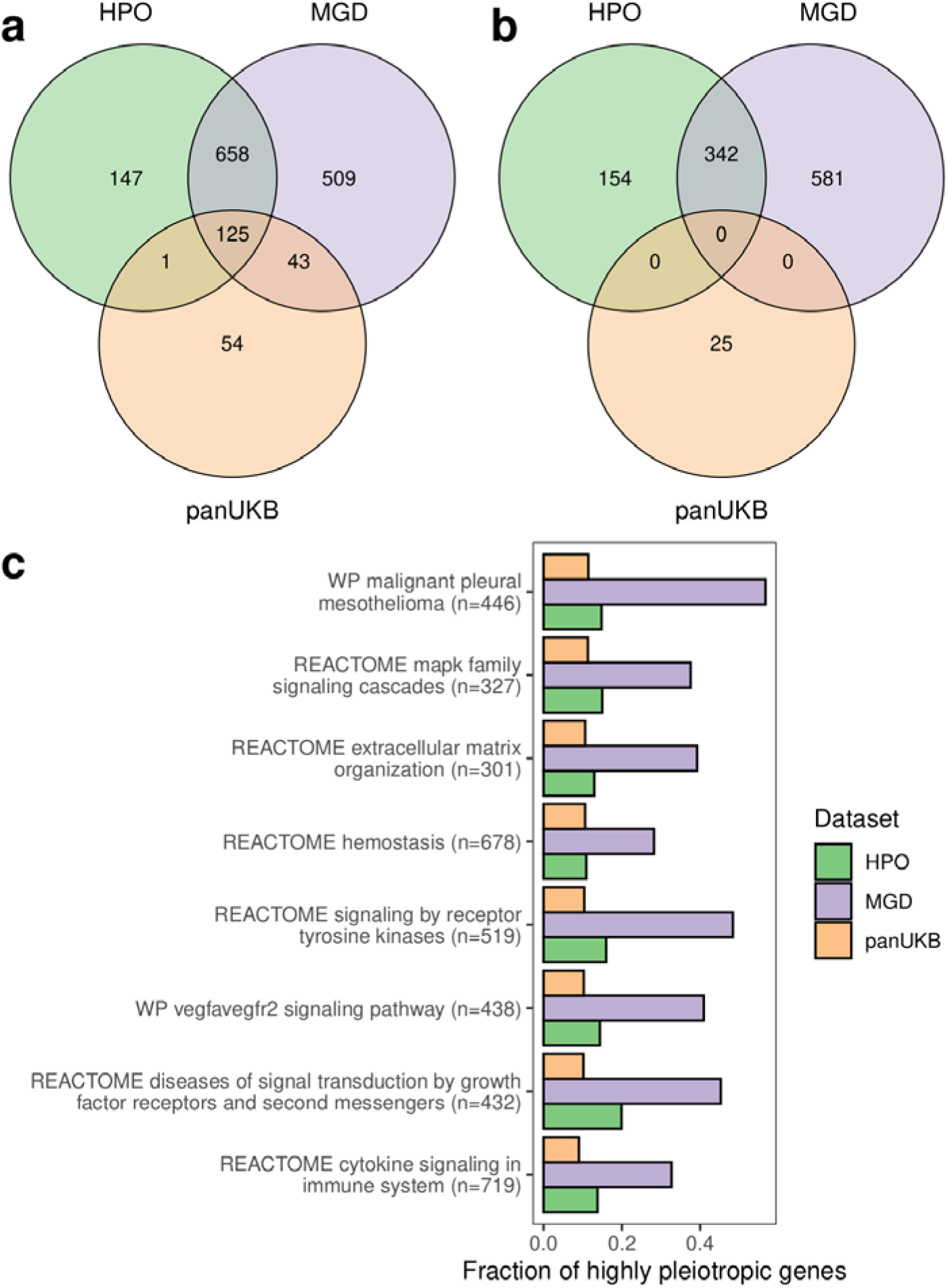
Pleiotropic genes in complex and Mendelian traits are involved in many common processes. (a, b) Venn diagrams showing the overlap between enriched GO terms identified using the highly pleiotropic gene sets from indicated data sources. All (a) or dataset-specific (b) highly pleiotropic genes were used for analysis. (c) A bar plot showing the fraction of highly pleiotropic genes for the 8 large canonical pathways that are significantly enriched with highly pleiotropic genes from all three datasets.

We next questioned if the observed overlap in the sets of enriched gene sets is driven by genes that are pleiotropic in both trait domains. To answer this question, we performed gene set enrichment analysis on highly pleiotropic GWAS genes that had no phenotype associations in either HPO or MGD. Importantly, such an analysis identified much fewer hits (25 canonical pathways, 8 GO BP terms, and 1 GO MF term). All 25 of these pathways and 7 out of 8 GO:BP terms were specific to GWAS data (that is, were not identified when using Mendelian-trait specific genes for gene set enrichment testing) (Figure 3b, Extended Data Figure 11). This result implies that, while there is a very substantial overlap in the set of biological processes controlled by highly pleiotropic genes from Mendelian and complex trait domains, this overlap is likely driven by genes that demonstrate pleiotropic effects in both domains. Domain-specific pleiotropic genes, on the other hand, tend to be involved in a specific set of processes. For the complex trait domain, this set of processes predominantly included various pathways associated with xenobiotic and steroid metabolism.

Having demonstrated the similarity in the sets of processes controlled by pleiotropic genes, we next went on to identify the biological processes, cellular components, and molecular function that are associated with the greatest proportion of highly pleiotropic genes across all datasets. To this end, we focused our analysis on a limited set of GO terms and pathways with large numbers of genes (see Methods for details), and then prioritized these gene sets by the minimum proportion of pleiotropic genes across datasets. Eight out of 56 pre-selected canonical pathways showed significant enrichment of highly pleiotropic genes in all three datasets. In good concordance with results described above, these pathways corresponded to cell proliferation-controlling signaling cascades (e.g., MAPK signaling cascade), genes involved in extracellular matrix organization, hemostasis, and immune system function (Figure 3c). In line with these findings, the highest proportion of highly pleiotropic genes was found for GO BP terms related to development, immune cell differentiation and activity, hemostasis, and response to extracellular stimuli (Extended Data Figure 12a). Within the cellular component ontology, pleiotropic genes from both trait domains were involved in transcriptional regulation complexes and chromatin, as well as in cell membrane regions and vesicles (Extended Data Figure 12b). In good concordance with these observations, molecular function terms linked to transcription factor binding, protein kinase binding or activity, and receptor binding, had the highest fraction of highly pleiotropic genes across all datasets (Extended Data Figure 12c). Of note, the proportions of genes with highly pleiotropic effects in mice were universally higher; thus, for the majority of the aforementioned terms, more than half of the corresponding genes had strong pleiotropic effects according to MGD data.

### Shared patterns of selection at pleiotropic loci across datasets

Results of the functional property analysis suggested that genes at pleiotropic loci (at least, within the monogenic trait domain) correspond to broadly expressed genes with large number of protein-protein interactions. This result predicts that these genes should experience greater pressure of negative natural selection constraining genetic variation within them. At the same time, it is not clear whether these patterns could be expected at pleiotropic GWAS loci. Hence, we next investigated the patterns of natural selection at pleiotropic genes and loci using several widely used measures of both negative (phastCons scores, loss-of-function variation and missense variation statistics from gnomAD) and positive (population-specific integrated Haplotype Scores (iHS) and the Density of Recent Coalescence events (DRC150)) selection).

When phastCons scores were used as a measure of negative selection, we found a significantly higher level of evolutionary conservation for highly pleiotropic genes in HPO and MGD (median gene-level score 0.66 and 0.68 compared to 0.63 and 0.64 for non-pleiotropic genes) (Figure 4a). For GWAS loci, no significant differences in gene-level phastCons scores were found. However, when the phastCons scores were averaged across a 100,000 base pair locus without taking genes into account, the pleiotropic loci show a weak enrichment with higher (> 0.15) phastCons scores (p = 0.017), though the median scores did not show any significant differences (Extended Data Figure 12a,d).

**Figure 4.**
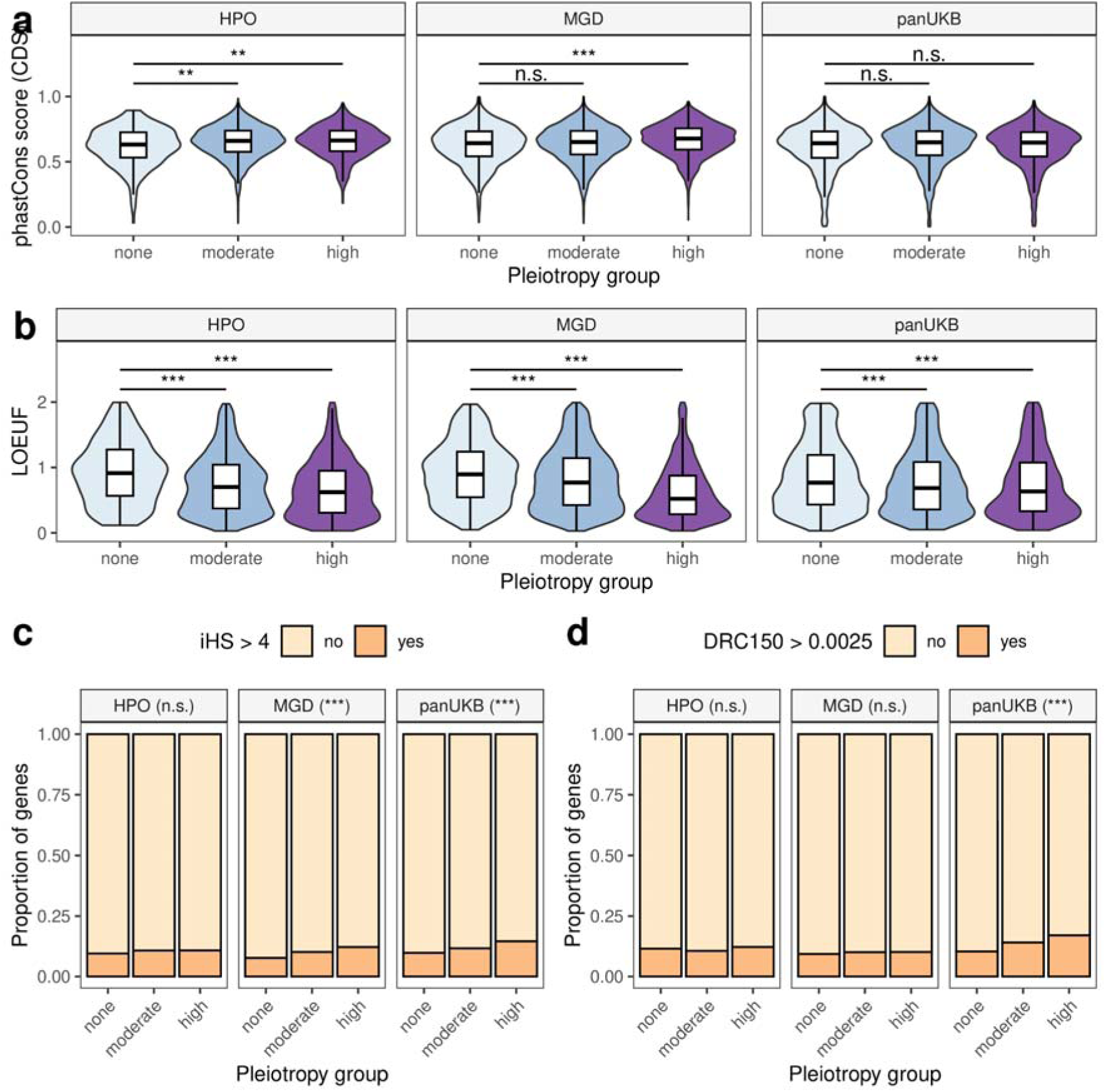
Pleiotropic genes are under greater evolutionary constraint despite bearing signals of recent adaptation. (a-b) Violin and box plots of the loss-of-function observed-to-expected upper fraction (LOEUF) (a) or gene-average phastCons score (b) values for genes in indicated pleiotropy groups. (c-d) Barplots showing the proportion of genes having maximum SNP iHS score > 4 (c) or density of recent coalescent events (DRC150) score > 0.0025 (d) in each pleiotropy group. ** - p < 0.01, *** - p < 0.001, n.s. - no significant differences in Wilcoxon-Mann-Whitney test (a, b) or chi-squared test (c, d) with Benjamini-Hochberg FDR adjustment. On (c, d) highly pleiotropic genes were compared to non-pleiotropic ones.

At the next stage of our analysis, we used gnomAD-derived gene-level constraint measures (LOEUF and pLI). Unlike phastCons scores, which are constructed using whole-genome alignments of multiple distantly related vertebrate species, LOEUF and pLI values reflect the loss-of-function (LoF) tolerance of human genes based on genetic variation within populations. With these measures being used, highly pleiotropic genes showed universally greater constraint (Figure 4b, Extended Data Figure 13a). Thus, the median LOEUF value for non-pleiotropic genes was 0.91, 0.90, and 0.77 for HPO, MGD, and pan-UKB data. For highly pleiotropic genes, the value dropped precipitously to 0.62, 0.52, and 0.63, respectively. A similarly pronounced difference was observed for the Z-score of the observed-to-expected missense variant ratio (Extended Data Figure 13b). When genes were split into categories according to their probability of loss-of-function intolerance (pLI) value, highly pleiotropic genes showed a universal and highly significant enrichment with LoF-intolerant (pLI > 0.9) genes across all data sources (30%, 33%, and 26% of highly pleiotropic genes were LoF-intolerant compared to 15%, 13%, and 19% of non-pleiotropic genes for HPO, MGD, and pan-UKB data; p << 0.001 in chi-squared test in all cases). Taken together, these results indicate that pleiotropic genes are under greater negative selection pressure in both complex and Mendelian trait domains.

We next went on to test the enrichment of pleiotropic genes and loci with recent positive selection signals. As mentioned earlier, two different measures of recent positive selection which capture different timescales of selection were utilized. As recent positive selection is not expected to operate uniformly on all genes in a group or across the entire gene sequence, we focused on comparing the proportions of genes for which the maximum value of proximal SNPs exceeded a threshold (a cutoff value of 4 was chosen for iHS, and 0.0025 - for DRC150). As can be seen in Figures 4c-d, we observed a general tendency towards the enrichment of pleiotropic genes with signals of recent positive selection, though the strength of this trend varied between the datasets and metrics. For instance, 12.2% of highly pleiotropic murine gene orthologs neighbored a variant with an iHS > 4, compared to 7.6% of non-pleiotropic ones (Figure 4c). Similarly, as many as 14.5% of highly pleiotropic genes in the pan-UKB GWAS data bore a recent positive selection signal according to iHS values, compared to only 9.7% of non-pleiotropic genes. The trend, however, was observed only with a cutoff of iHS > 4, and was accompanied by only a subtle shift in the median value (2.44 vs. 2.12, Extended Data Figure 14). Moreover, comparison of maximum iHS value for each locus (irrespective of neighboring genes) did not show any significant difference in the iHS value for different pleiotropy groups (Extended Data Figure 12b,e). When the DRC150 value was used as measure of positive selection, only highly pleiotropic GWAS genes showed a significant enrichment with large DRC150 values (17.0% compared to 10.3% for non-pleiotropic genes, according to closest gene-based data, Figure 4d). Unlike that observed with iHS, the effect of pleiotropy on DRC150 was by far more pronounced when a maximum value of DRC150 for all variants in a 100,000 bp locus was evaluated (33.9% vs 10.5%, Extended Data Figure 12c,f).

Given the above observations, we next questioned if highly pleiotropic loci are enriched with genes that bear a combination of negative and positive selection signatures. To answer this question, we selected a subset of human genes with strong selective constraint against loss-of-function variation (LOEUF < 0.6) and a nearby signal of recent positive selection (an SNP with iHS > 4). A total of 730 (5.3%) of all genes met these conditions. The proportion of such genes among non-pleiotropic ones was similar (4.2% in HPO, 3.4% - in MGD, and 3.2% - in GWAS). Among the highly pleiotropic genes, however, this proportion was significantly higher for MGD and pan-UKB data (8.1% and 7.0%, respectively) (Extended Data Figure 15a). Similar results were also observed when using a combination of low LOEUF and high DRC150 value (Extended Data Figure 15b).

Finally, we asked if the observed overlap between signals of negative and positive selection can be explained by the general correlation between gene-level constraint (LOEUF) and iHS/DRC150 that may be driven by background selection. To answer this question, we compared the LOEUF values for genes with and without a neighboring positive selection signal (according to iHS or DRC150). This analysis showed that, while genes bearing an SNP with iHS > 4 indeed tend to be under greater selective constraint, an inverse tendency is observed for DRC150 (Extended Data Figure 16). Hence, we conclude that the observed enrichment of overlapping signals of negative and positive selection can not be explained solely by the effects of background selection.

Taken together, these findings demonstrate a universal propensity of pleiotropic genes to be under increased selective constraint, and validate earlier suggestions regarding a possible enrichment of pleiotropic loci with positive selection signals. Moreover, our results reveal a unique enrichment of pleiotropic genes with those bearing combined signals of positive and negative selection.

## Discussion

The phenomenon of pleiotropy has attracted the attention of researchers since the early years of genetics. It is now well-established that pleiotropy is abundant in natural systems, and the distribution of the degree of pleiotropy usually has a characteristic L-shape (reviewed in Zhang, 2023). All three data sources used in our analysis also showed a characteristic left skew of the distribution; however, we observed substantial differences in the median degree of pleiotropy as well as the estimated modularity of the underlying genotype-to-phenotype networks. Thus, gene-level pleiotropy estimated using HPO and MGD data was much higher compared to the genome-wide association data. Earlier studies based on mouse phenotype data also showed a higher median degree of pleiotropy (Wang et al., 2010; Munoz-Fuentes et al., 2022). It is necessary to consider the possible sources of these differences.

The most natural explanation of the results is the different nature of traits and mutations used to construct the data. For HPO, the main source of gene annotation is the phenotype observed in patients with Mendelian disease caused by a pathogenic variant in a given gene, usually being a loss-of-function allele (Kohler et al., 2021). Similarly, phenotype data for mice comes from single gene or high-throughput knockout assays. As thus, genotype:phenotype relationships recorded in both HPO and MGD come from high impact genetic variants which are commonly assumed to have higher degree of pleiotropy compared to regulatory variation (Carroll, 2005). On the other hand, genetic variation used to conduct genome-wide association analysis is mostly regulatory or silent, though pleiotropic variants are known to be enriched for missense variants (Watanabe et al., 2019; Shikov et al., 2020).

While the trait and mutation type could explain the observed differences in the degree of pleiotropy, it is complicated to rule out the possibility of a technical confounding. For example, upper-level Mammalian Phenotype ontology terms are not entirely independent, with possible implications for pleiotropy analysis (Munoz-Fuentes et al., 2022). In our analysis, we chose upper-level MP terms over other ontology term-based metrics as they maximized the number of non-pleiotropic genes and minimized the median degree of pleiotropy (see Supplementary Note, section 1), However, we believe that the phenotype definition still plays a role in inflating the degree of pleiotropy in certain cases.

For genome-wide association data, on the other hand, the estimate of the gene-level degree of pleiotropy could be impacted by the causal gene selection method based on summary statistics (see Supplementary Note section 2), as well as method for independent phenotype definition (Supplementary Note section 3). In our work, we’ve used phenotypic correlation-based clustering to reduce the impact of vertical pleiotropy, as we’ve previously shown a positive impact of this procedure on the phenome-wide analysis results (Shikov et al., 2020). While this procedure efficiently reduces the observed degree of pleiotropy, it is still possible that many complex traits with relatively low observed phenotypic correlation could still fall under the same upper-level phenotype ontology term. Hence, we could expect that mapping complex traits to phenotype ontology terms could further reduce the observed statistical pleiotropy in GWAS data.

In our study, we showed that genes bearing highly pleiotropic variation are more broadly expressed (Figure 2a, Extended Data Figure 2), have more protein-protein interactions (Figure 2b) and a richer set of molecular functions (Extended Data Figure 3), participate in more biological processes (Figure 2c), are more dosage sensitive (Figure 2d, Extended Data Figure 4), and, finally, are under greater selective constraint (Figure 4a-b, Extended Data Figure 13). These results are in line with earlier observations made in model organisms (He and Zhang, 2006) and in human complex traits (Shikov et al., 2020). A more intriguing finding, however, is the striking similarity in the properties of highly pleiotropic genes in complex traits and Mendelian traits, which is observed despite the aforementioned discrepancies in the degree of pleiotropy. Besides the same direction of differences between non-pleiotropic and highly pleiotropic genes across datasets, it is also important to emphasize that a highly pleiotropic gene according to GWAS data was more similar to a moderately pleiotropic or non-pleiotropic gene according to HPO or MGD data. These observations are in line with a lower median degree of pleiotropy observed in the genome-wide association data. Nevertheless, we can conclude that the same mechanisms guide pleiotropic effects in both complex and Mendelian trait domains.

It is also worth noting that the differences in all gene-level metrics tested were usually much more pronounced in MGD than in HPO data, and certain trends (e.g., the enrichment of highly pleiotropic genes with recent positive selection signals) were significant in MGD, but not HPO. These observations suggest that the phenotype of high-impact genetic variation is much better described in mice than in humans. Indeed, large-scale phenotyping efforts such as the International Mouse Phenotype Consortium (IMPC) (Meehan et al., 2017; Cacheiro et al., 2019) have greatly enhanced the depth of mouse phenotype description. In humans, on the other hand, description of the phenotype is only performed by examining the patients with rare disease. Hence, many genes may lack phenotype annotation due to various factors, such as survival of the patient until the time of molecular diagnostics or the ability to clearly demonstrate the causal effect of a genetic variant observed in a patient.

Besides shared properties of pleiotropic genes, our analysis identified a set of processes that are enriched with pleiotropic genes across species and trait domains. This set of pathways includes ones involved in cell proliferation, signaling, immune system function, and extracellular matrix organization (Figure 3, Extended Data Figure 11). These findings are in good concordance with our earlier analysis of pleiotropic gene functions in complex traits (Shikov et al., 2020). Besides, the present analysis indicates that the previously observed enrichment of highly pleiotropic loci with liver-specific genes is the unique feature of complex trait domain. Thus, xenobiotic metabolism and related pathways were enriched among GWAS-specific highly pleiotropic genes, but not among highly pleiotropic Mendelian trait genes in either species.

Shared functions of highly pleiotropic genes across trait domains (Figure 3, Extended Data Figure 10) suggest that a highly similar set of pathways is responsible for pleiotropic effects of genetic variation; however, the magnitude of these effects would be dependent on the type of mutation (e.g., null (loss-of-function) mutation, change in the amino acid sequence of a protein, or a variant in the regulatory element) (Figure 5). Thus, high-impact genetic variants in crucial pathway genes would have pronounced pleiotropic effects and manifest in Mendelian phenotypes (rare congenital disorders), and would be subject to enhanced purifying selection. Lower-impact variants in the same genes would also have pleiotropic effects, albeit with a lower magnitude, driving multiple complex traits rather than Mendelian phenotypes. Such variants will be enriched in adaptive alleles, in accordance with both our (Figure 4) and earlier findings (e.g., Frachon et al., 2017; Sakaue et al, 2021). In pathways with a restricted set of functions, on the other hand, all types of mutations would likely lack pleiotropic effects, and would hence be both less adaptive and subject to lower negative selective pressure.

**Figure 5.**
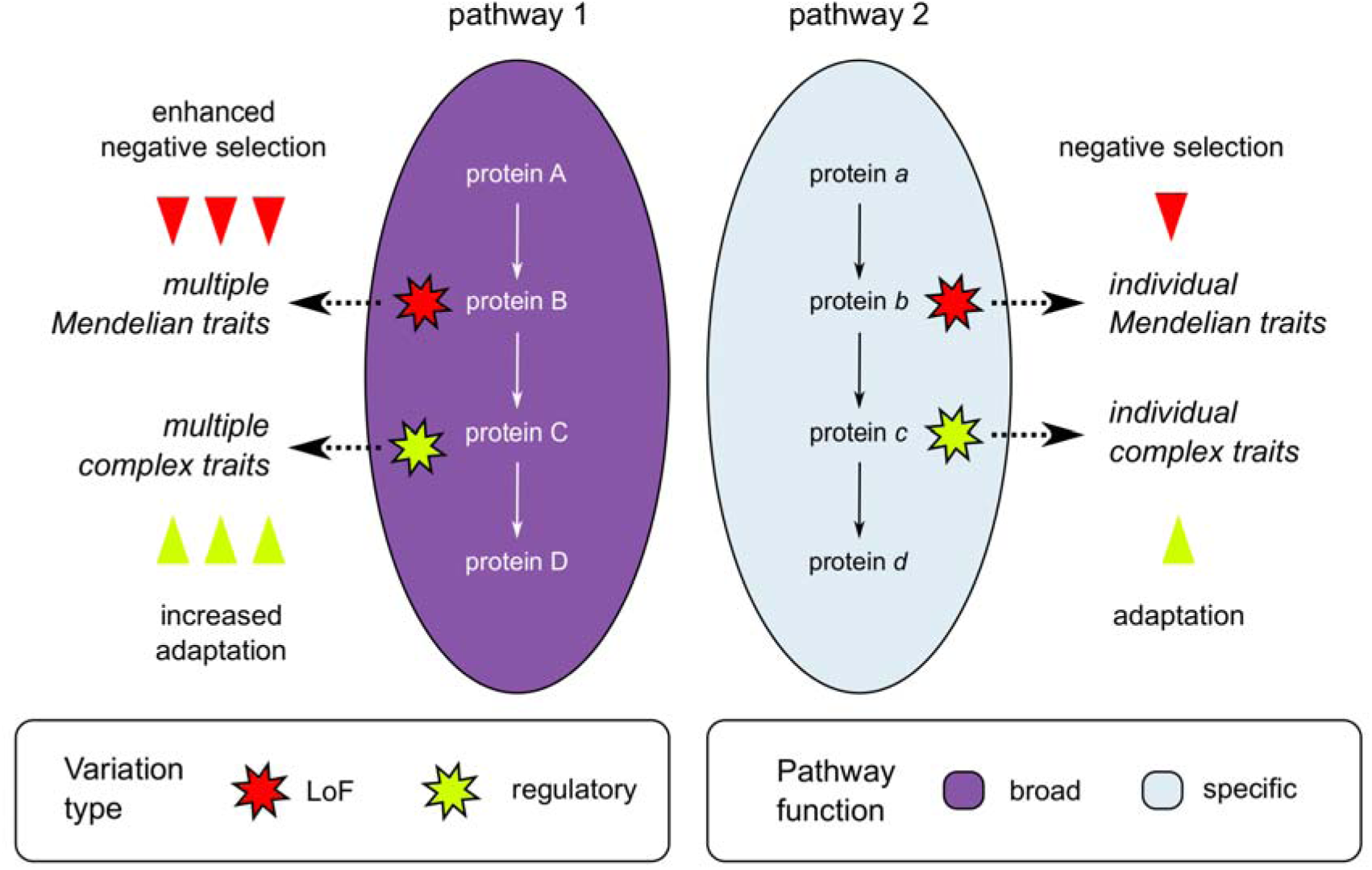
A model for pleiotropic effects in different trait domains. Ellipses represent pathways colored by their functional relevance in an organism (i.e., broad or tissue/cell type-specific), arrows connecting proteins within pathways represent the direction of signal transduction or position of genes in metabolic reaction chains. Polygonal stars represent mutations that are colored according to their type (LoF or regulatory) See text for further explanation of the model.

In our previous work, we observed an interesting tendency of pleiotropic genetic variants in GWAS to have higher allele frequency (Shikov et al., 2020). At the time, we have proposed three explanations for this observation: (i) removal of rare variants at pleiotropic loci due to strong purifying selection; (ii) greater power to detect genome-wide associations for common genetic variants; and (iii) spurious pleiotropy of common variants that tag different rare causal variants at the same locus. In addition to the three hypotheses mentioned above, one might also suggest that higher frequency of pleiotropic variants may be driven by an enrichment of loci with recent positive selection signals, an assumption that has been supported by evidence in some recent studies (Frachon et al., 2017; Sakaue et al., 2021). The results included in the present study indicate that pleiotropy correlates both with increased strength of purifying selection and higher frequency of adaptive alleles, favoring both the first and fourth hypotheses. Moreover, our results show that both negative selection strength and the enrichment with positive selection signals is highest for highly pleiotropic genes. This is especially true for genes with pleiotropic effects in mice, which showed both the strongest negative selection against loss-of-function variants and the most pronounced enrichment with adaptive alleles (according to the iHS value). These observations contradict earlier findings and predictions that pleiotropy should only increase the rate of adaptation for moderate degrees of pleiotropy (Wang et al., 2010; Frachon et al., 2017). It is important to note, however, that the enrichment with more recent positive selection (detected using the DRC150 statistic), was only observed in GWAS data, where the cutoff for selection of highly pleiotropic genes was lower. Still, the proportion of genes with high DRC150 value increased when higher degree of pleiotropy was demanded (e.g., 32 out of 168 genes had a positive selection signal when requiring at least 8 trait clusters). Hence, we believe that our results emphasize that a high degree of pleiotropy is, at the very least, compatible with active positive selection.

To sum up, pleiotropy is not only a basic feature of genotype-to-phenotype networks, but an inevitable consequence of the organismal complexity. Still, while pleiotropy imposes additional constraint on genetic variation, it appears to provide additional opportunities for adaptation of complex organisms to changing environments.

## Materials and methods

### Genotype-to-phenotype data collection

To compile a dataset consisting of genotype-to-phenotype relationships in the Mendelian trait domain, we obtained information about orthologous gene pairs between mice and humans from the Mouse Genome Database (MGD) (the HMD_HumanPhenotype file was used; https://www.informatics.jax.org/downloads/reports/HMD_HumanPhenotype.rpt). For each ortholog, we included associations with phenotype terms from the Mammalian Phenotype ontology (MP) for mice and the Human Phenotype Ontology (HPO) for humans. For human genes, gene annotations were downloaded directly from the HPO website (https://hpo.jax.org/data/annotations, accessed on 2024-07-01). For MP data, information was retrieved from the MGI_PhenoGenoMP file, https://www.informatics.jax.org/downloads/reports/MGI_PhenoGenoMP.rpt).

As a source of genotype-to-phenotype relationship information in complex trait domains, we utilized publicly available summary statistics of genome-wide association analysis by the pan-UK Biobank study (Karczewski et al., 2024). The pan-UKB manifest file with heritability estimates was pre-processed to select traits with *P(h^2^* = 0*) <* 0.01 according to the standard normal distribution (Z-score > 2.32). For traits with different encoding and pre-processing methods used in the original pan-UKB study, a single entry with the maximum heritability estimate was selected. Summary statistics files were downloaded for the resulting set of 1263 traits (Supplementary Table S1).

### Definition of pleiotropy in phenotype ontology data

To calculate the degree of pleiotropy of a gene, all phenotype ontology terms associated with a particular gene were converted to upper-level terms using the ontobio package (for HPO data, terms were converted to MP to simplify further cross-species comparison, official upper-level term mapping provided by MGI was used (https://github.com/mgijax/mammalian-phenotype-ontology/tree/main/mappings)). Then, the number of such upper-level MP terms linked to each gene was used as a measure of its degree of pleiotropy. To identify highly pleiotropic genes, we split all genes into quartiles according to their degree of pleiotropy and then considered all genes in the fourth quartile as highly pleiotropic ones. Besides the aforementioned approach, clustering of all phenotype ontology terms was attempted as a measure of pleiotropy, with results of the analysis being similar to those obtained with upper-level terms (see Supplementary Note, section 1 for details).

### Definition of pleiotropy in genome-wide association data

Summary statistics of pan-UKB GWAS were used to identify lead SNPs using the clumping procedure available in PLINK v. 1.9 (Purcell et al., 2007) and the 1000 Genomes phase 3 genotype data (Auton et al., 2015). Following clumping, the closest gene was retrieved for each of the identified lead SNPs using BEDtools (Quinlan and Hall, 2010) and the GENCODE v19 human gene annotation file (Frankish et al., 2020). An alternative approach based on all genes within a 100,000 bp interval around each lead SNP was also attempted, with similar results (see Supplementary Note, section 2).

A matrix of gene-trait associations was constructed using the inferred causal gene information. Results of hierarchical trait clustering using the phenotypic correlation estimates provided by the authors of the pan-UKB study were used to merge associations by cluster (for each cluster, a gene was considered to be associated with the cluster if it was identified as the closest gene for a lead SNP in GWAS results for at least one trait). The number of associated trait clusters was used as the resulting measure of the degree of pleiotropy, similar to our earlier work (Shikov et al., 2020). Similarly to the analysis of phenotype ontology terms, top 25% of all genes with the highest degree of pleiotropy were considered as highly pleiotropic, and all other genes with the degree of pleiotropy greater than 1 were considered as moderately pleiotropic.

### Analysis of genotype-to-phenotype network modularity

To perform an analysis of modularity of the bipartite gene-trait network, preprocessed gene-to-trait associations from each data source were converted into an incidence matrix. The matrix was then used to construct a bipartite graph using the igraph package (Casardi et al., 2006), followed by greedy community detection and calculation of network modularity score.

### Comparison of functional gene properties

Summary statistics of gene expression profile (median expression across tissues, number of tissues with expression at >= 5 TPM, maximum expression across tissues) were calculated using the Genotype Tissue Expression (GTEx) v8 data (Lonsdale et al., 2013). The number of protein-protein interactions recorded for a gene was retrieved using the BioGRID database (Oughtred, et al., 2021). Additionally, gene-level probabilities of dosage sensitivity (pHaplo and pTriplo) were taken from a paper by Collins et al., 2022 (Collins et al., 2022). Finally, the number of Gene Ontology (GO) (Ashburner et al., 2000) terms associated with each gene was calculated using the org.Hs.eg.db and org.Mm.eg.db packages for R.

Besides the analysis of numeric features, a gene set enrichment analysis using the GO term enrichment was performed for highly pleiotropic genes using the clusterProfiler package for R (Yu et al., 2012). In addition to GO terms, hallmark gene sets and canonical pathways (CP) from the Molecular Signatures Database (MSigDB) (Liberzon et al., 2011; Liberzon et al., 2015) were used in gene set enrichment analysis.

For the analysis of gene sets with the most consistent enrichment across datasets, we merged the results of enrichment analysis for the three data sources into a single dataset (per each gene set group (hallmarks, canonical pathways, and BP, CC, and MF branches of GO), and then ordered the gene sets by the minimum percentage of highly pleiotropic genes across the data sources. For GO terms, only large (> 500 genes) terms were considered. For a largely redundant CP collection, a set of mostly non-overlapping pathways was constructed as follows: (1) all pathways were ordered by the number of genes; (2) the largest pathway was selected, and all genes corresponding to this pathway were removed; (iii) the procedure was repeated with the remaining genes until at least one pathway with > 25 genes remained. This procedure resulted in a set of 56 (out of initial 2,982) mostly non-redundant pathways covering more than 87% (11,681/13,343) of all human genes with known function (Supplementary Table S2).

### Signals of positive and negative selection

For the analysis of purifying selection for pleiotropic and non-pleiotropic loci, gene-level constraint measures for human genes were retrieved from the Genome Aggregation Database (gnomAD) v.2.1. (Karczewski et al., 2020). Loss-of-function variants observed-to-expected ratio upper fraction (LOEUF), Z-score for observed-to-expected missense variant ratio, and the probability of loss-of-function intolerance (pLI) (Lek et al., 2016) were used for gene annotation and comparison. Additionally, we used phastCons (Siepel et al., 2005; 10.1101/gr.3715005) conservation scores computed using 100 vertebrate species obtained from the UCSC Table Browser (https://genome.ucsc.edu/cgi-bin/hgTables, accessed on 10 July 2024). The phastCons scores in bigWig format were converted to BEDGRAPH format using the bigWigToBedGraph tool (https://github.com/ENCODE-DCC/kentUtils, accessed on 20 July 2024), and then averaged over the gene or locus interval.

To assess the overlap with recent positive selection signals, two main metrics were employed that capture different time periods: (i) the integrated haplotype score (iHS) and (ii) density of recent (over the last 150 generations) coalescence events (DRC150). The iHS values for the British (GBR) subpopulation from 1000 Genomes project (Auton et al., 2015) were taken from a recent analysis by Johnson et al. (Johnson et al., 2018). DRC150 were taken from the original study by Palamara et al. (Palamara et al., 2018). For both metrics, gene-wise iHS and DRC150 value was calculated by intersecting the gene interval (for HPO/MP data) or the 100,000 bp interval around the lead SNP (for the pan-UKB data) with SNP coordinates (annotated with respective iHS and DRC150 values), and the maximum absolute value of each metric was retrieved for a gene or locus in question.

## Data availability

All data and code pertinent to the analysis presented in this work are available through GitHub: https://github.com/ibre-research/pleiotropy_analysis/.

## Acknowledgements

We thank Systems Biology Fellowship for providing support for Y.A.B. in 2020 - 2022. We are grateful to Anton I. Changalidis for his help in acquisition and curation of the pan-UKB dataset.

