## Extended Data Figures & Supplementary Note for "Functional determinants and evolutionary consequences of pleiotropy in complex and Mendelian traits"

[1] - Institute of Bioinformatics Research and Education, Belgrade Serbia
[2] - JetBrains Research, Belgrade, Serbia

### Extended Data

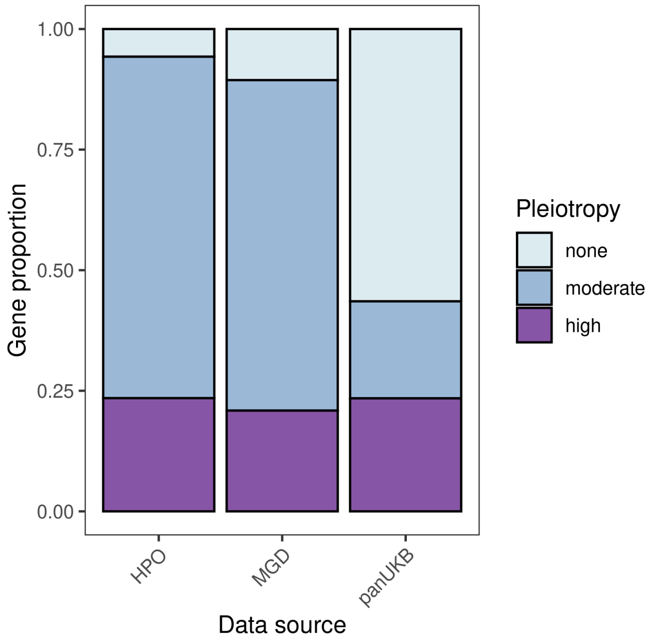


**Extended Data Figure 1. Proportion of genes in the three pleiotropy groups in each of the four datasets included in the analysis.** Note that the proportion of highly pleiotropic genes is almost constant due to the criteria used to identify highly pleiotropic genes (see Methods).

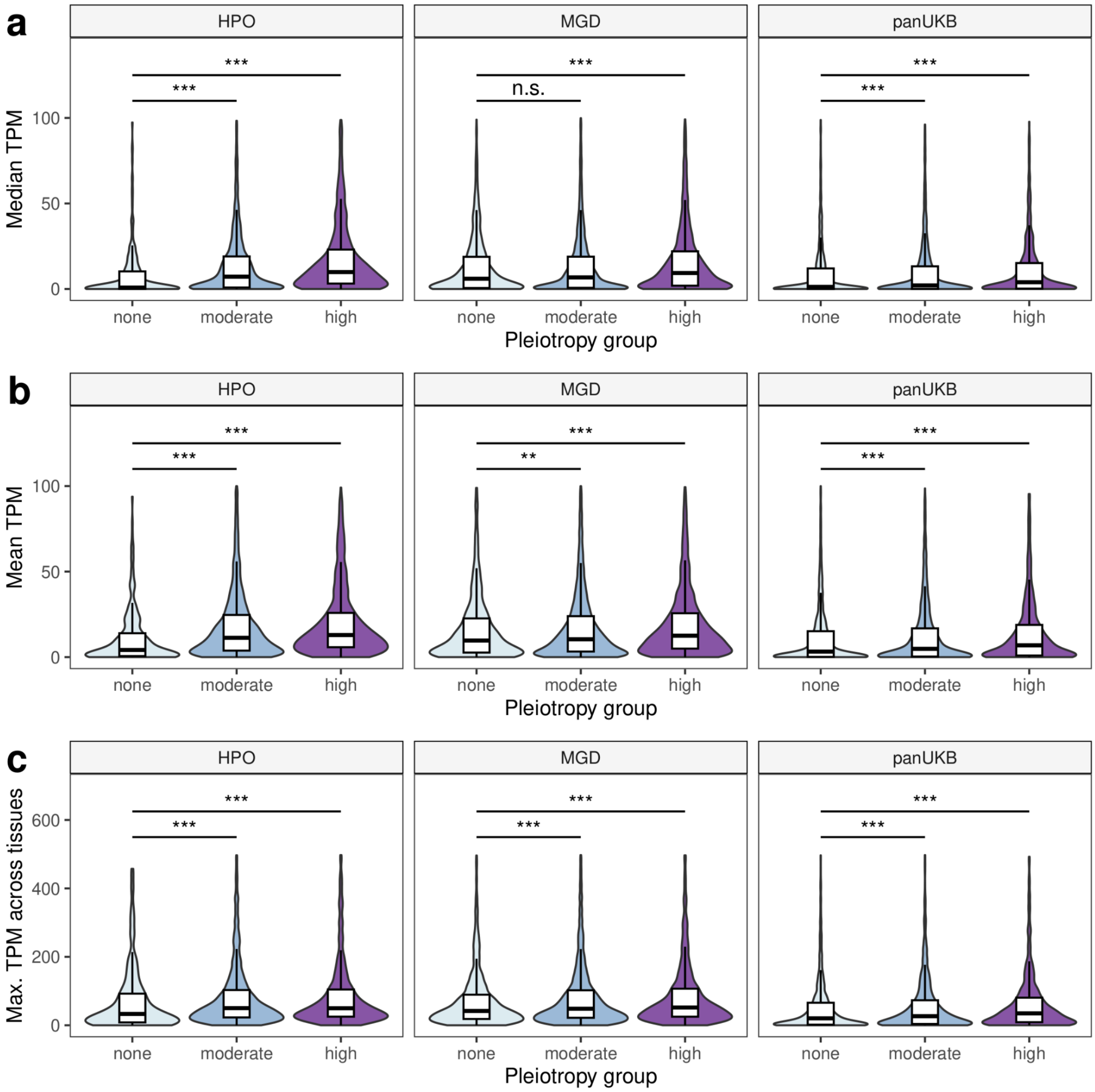


**Extended Data Figure 2. Pleiotropic genes have higher expression levels.** Shown are violin plots of the median (a), mean (b), and maximum (c) per-tissue expression level of the genes in indiciated pleiotropy groups. All expression values are in transcripts per million (TPM) according to the Genotype Tissue Expression (GTEx) data * - $p < 0.05$, ** - $p < 0.01$, *** - $p < 0.001$ in Wilcoxon-Mann-Whitney rank sum test with Benjamini-Hochberg FDR adjustment.

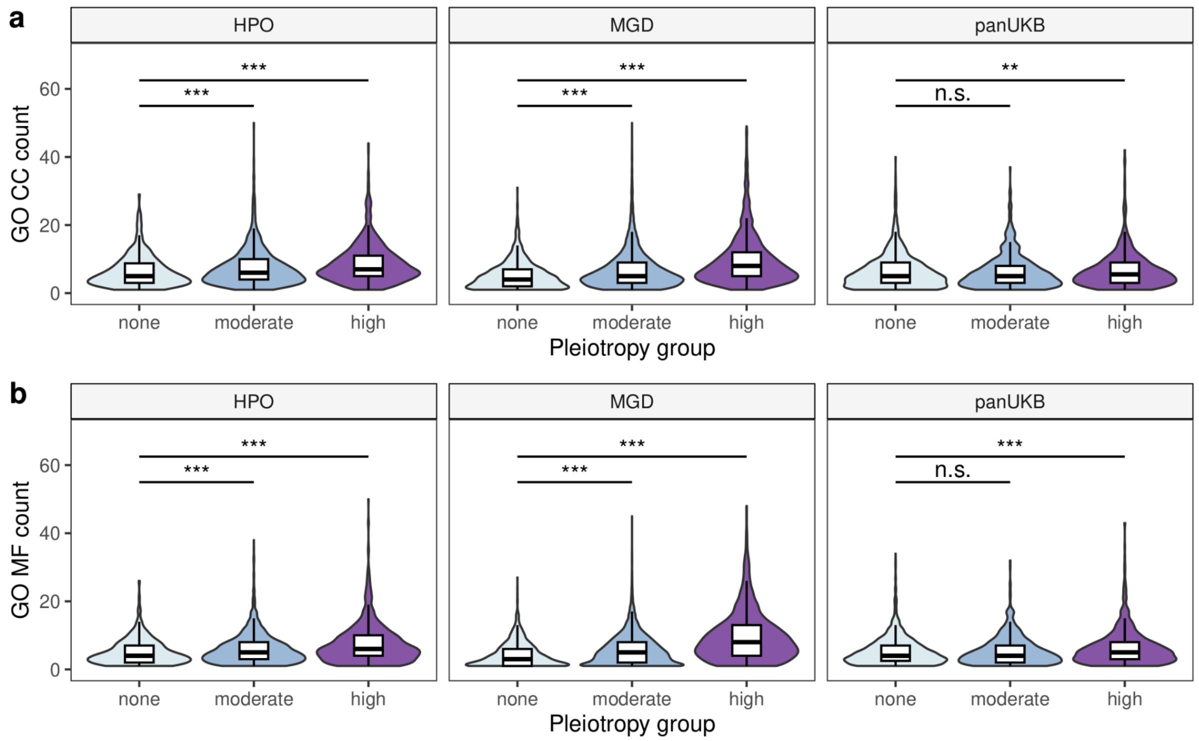


**Extended Data Figure 3. Highly pleiotropic genes have more biological functions.** Shown are violin plots of the number of cellular component (CC) (a) and molecular function (MF) (b) terms associated with genes in indiciated groups. * - $p < 0.05$, ** - $p < 0.01$, *** - $p < 0.001$ in Wilcoxon-Mann-Whitney rank sum test with Benjamini-Hochberg FDR adjustment.

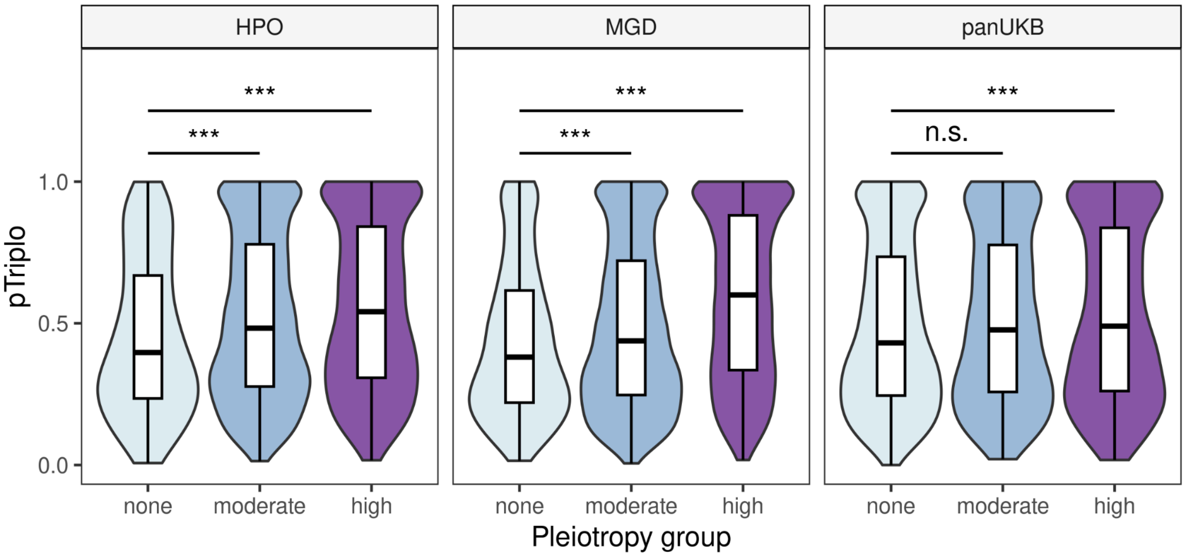


**Extended Data Figure 4. Highly pleiotropic genes have higher levels of triplosensitivity.** Shown are violin plots and box plots of the probability of triplosensitivity (pTriplo) according to data in Collin *et al*., 2022, ** - $p < 0.01$, *** - $p < 0.001$ in Wilcoxon-Mann-Whitney rank sum test with Benjamini-Hochberg FDR adjustment.

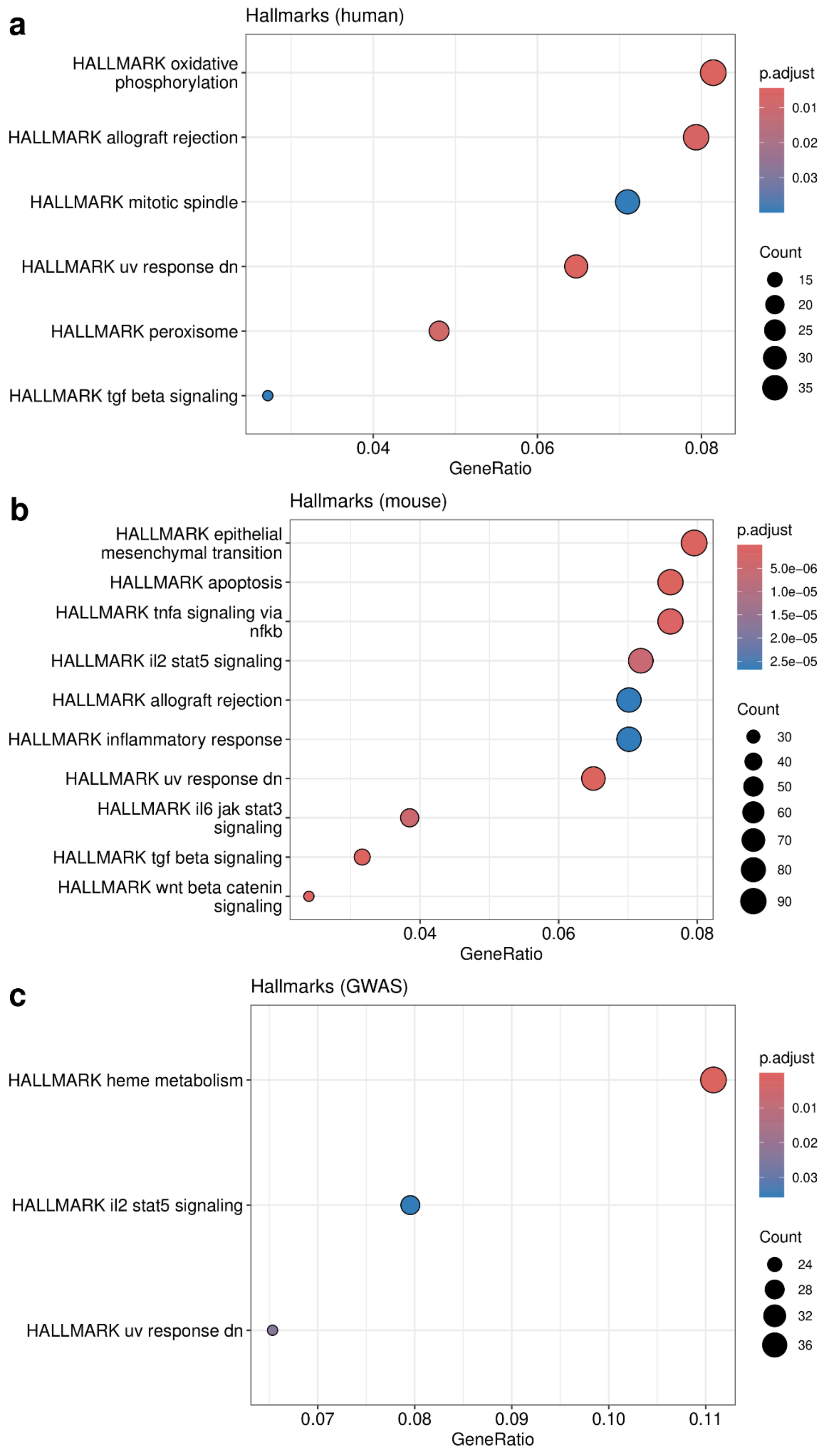


**Extended Data Figure 5. Hallmark gene sets associated with high degree of pleiotropy.** Shown are dot plots of the gene set enrichment analysis results of highly pleiotropic genes according to HPO (a), MGD (b), and pan-UK Biobank GWAS data (c). The analysis was performed against hallmark gene sets from MSigDB. Top 10 terms are shown in each case.

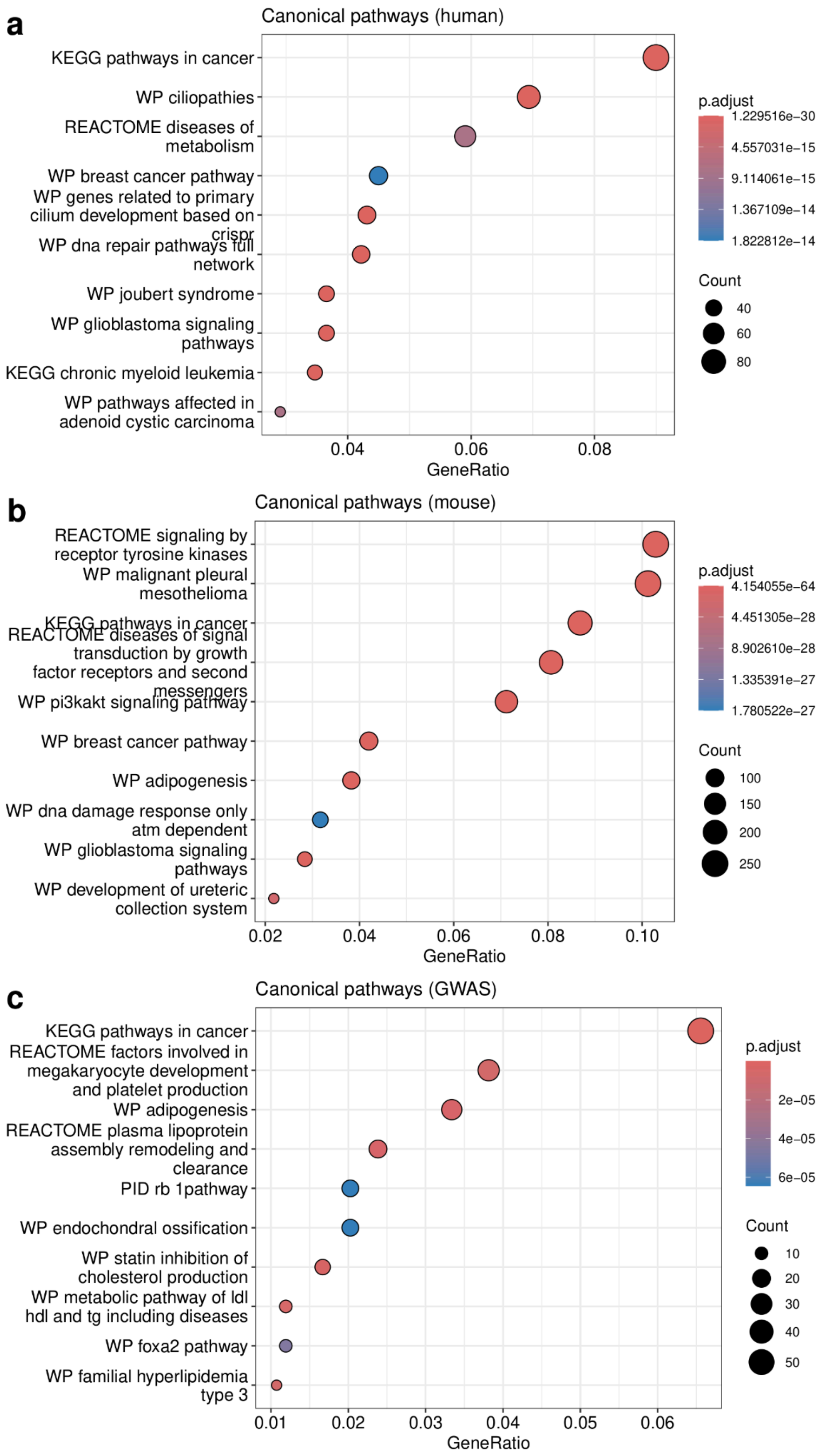


**Extended Data Figure 6. Canonical molecular pathways associated with high degree of pleiotropy.** Shown are dot plots of the gene set enrichment analysis results of highly pleiotropic genes according to HPO (a), MGD (b), and pan-UK Biobank GWAS data (c). The analysis was performed against canonical pathway (C2:CP) gene sets from MSigDB. Top 10 terms are shown in each case.

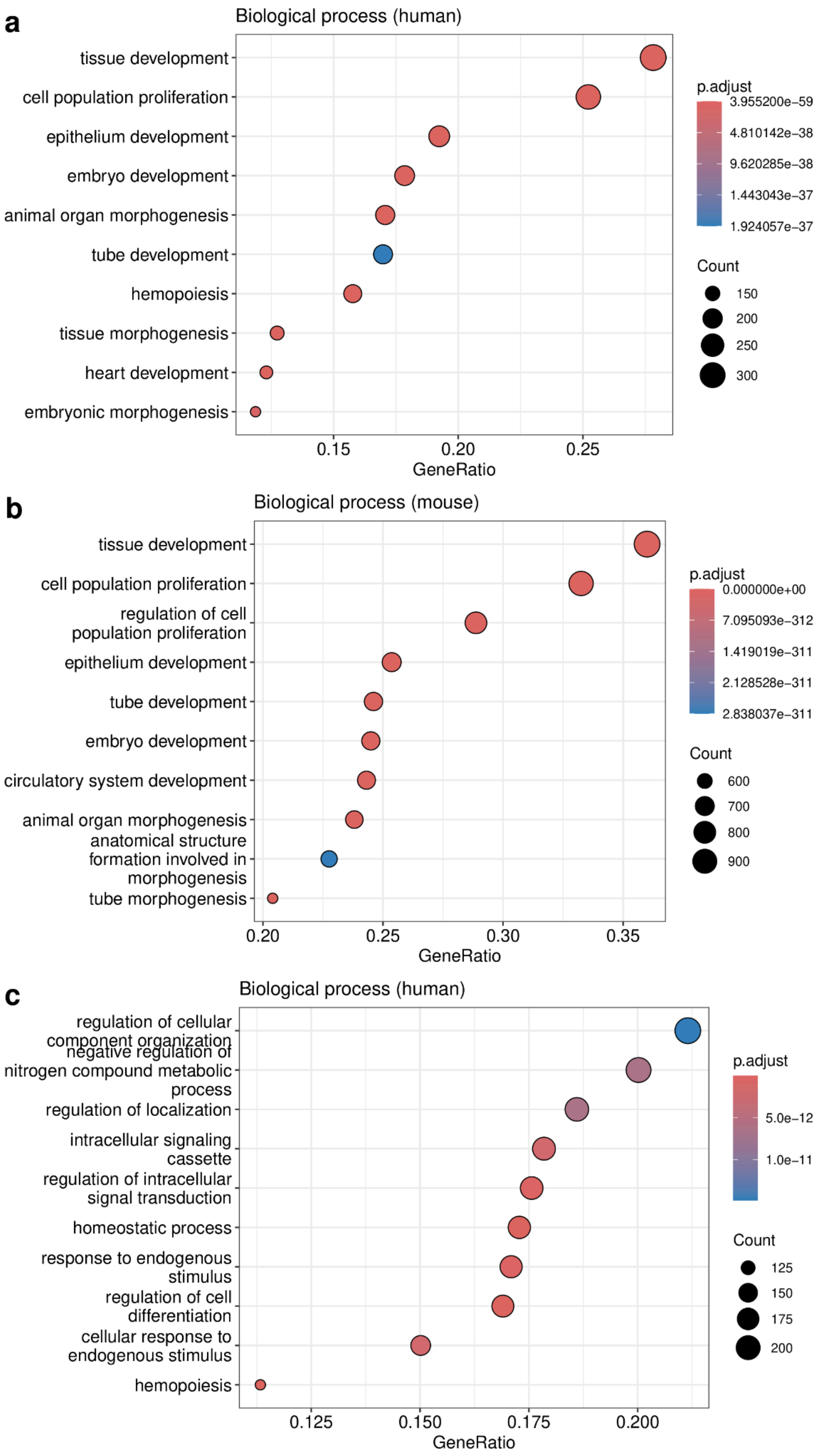


**Extended Data Figure 7. Biological processes associated with high degree of pleiotropy.** Shown are dot plots of the gene set enrichment analysis results of highly pleiotropic genes according to HPO (a), MGD (b), and pan-UK Biobank GWAS data (c). The analysis was performed against GO biological process terms. Top 10 terms are shown in each case.

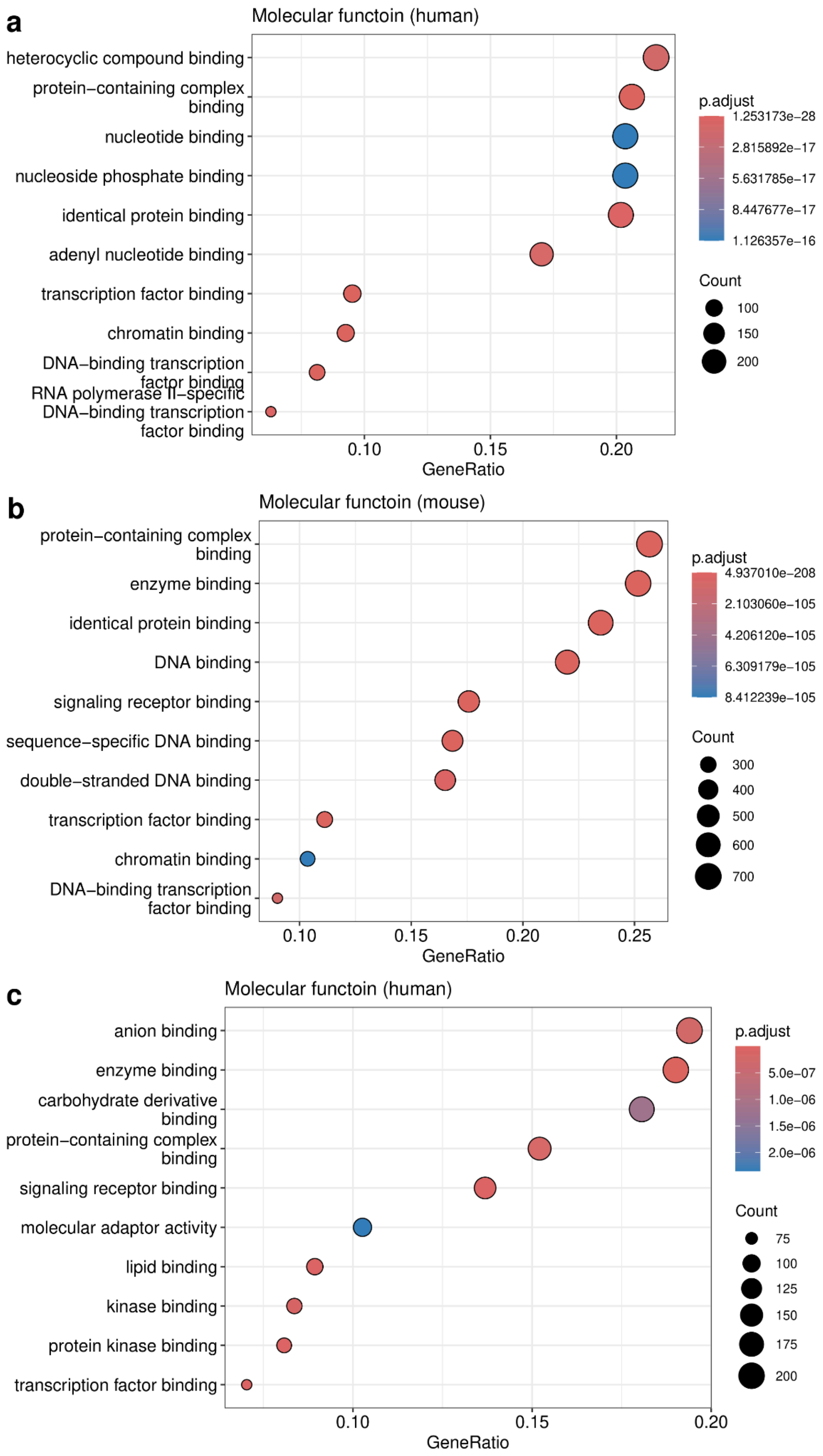


**Extended Data Figure 8. Molecular functions associated with high degree of pleiotropy.** Shown are dot plots of the gene set enrichment analysis results of highly pleiotropic genes according to HPO (a), MGD (b), and pan-UK Biobank GWAS data (c). The analysis was performed against GO molecular function terms. Top 10 terms are shown in each case.

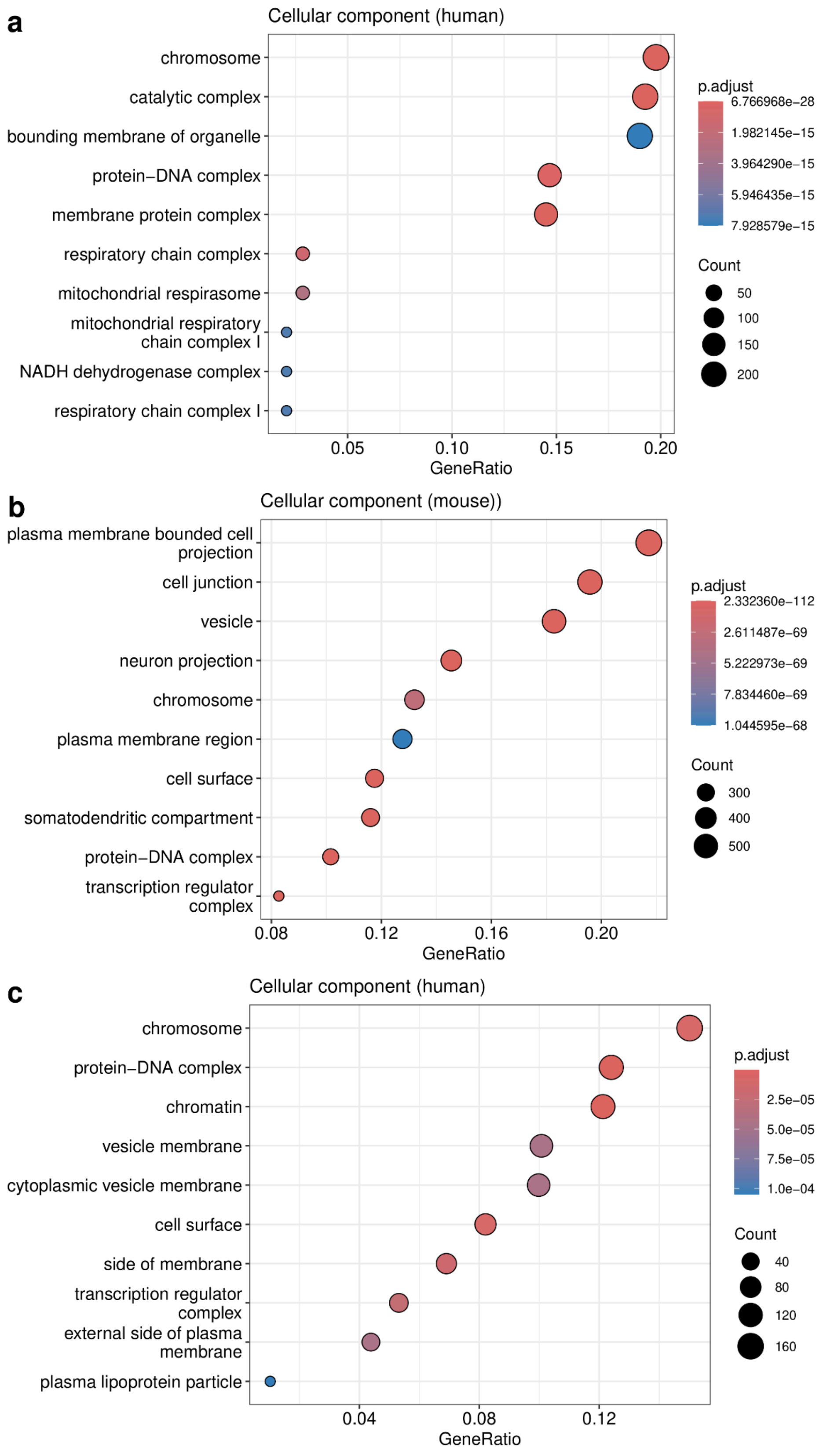


**Extended Data Figure 9. Cellular components associated with high degree of pleiotropy.** Shown are dot plots of the gene set enrichment analysis results of highly pleiotropic genes according to HPO (a), MGD (b), and pan-UK Biobank GWAS data (c). The analysis was performed against GO cellular component terms. Top 10 terms are shown in each case.

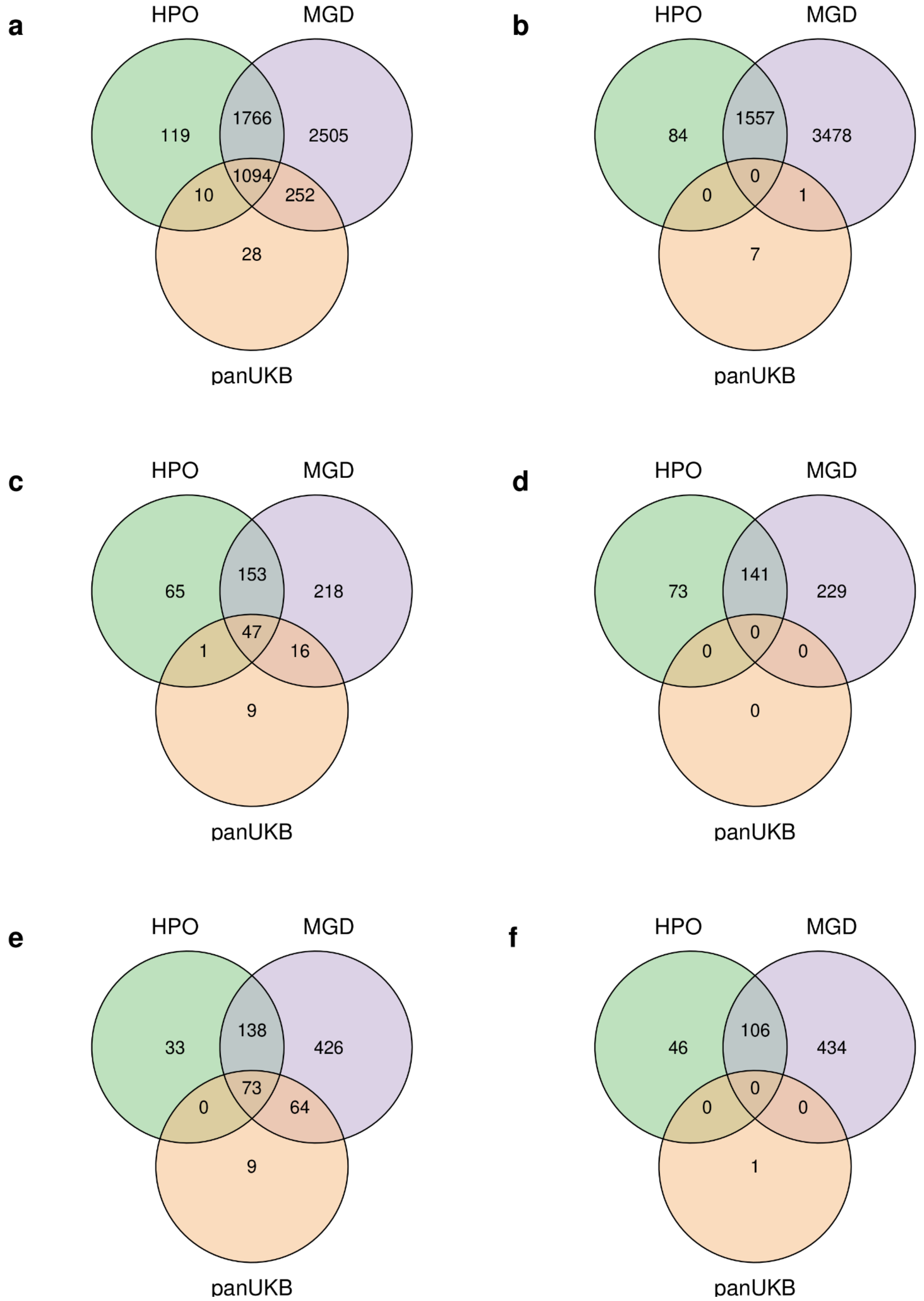


**Extended Data Figure 10. Highly pleiotropic genes share a large fraction of enriched GO terms.** Shown are Venn diagrams representing the overlap between sets of significantly enriched biological process (a-b), cellular component (c-d) and molecular function (e-f) terms for highly pleiotropic genes in the indicated datasets. On (b, d, f), genes having phenotype associations only in a given dataset (but not the other ones) were used for enrichment testing.

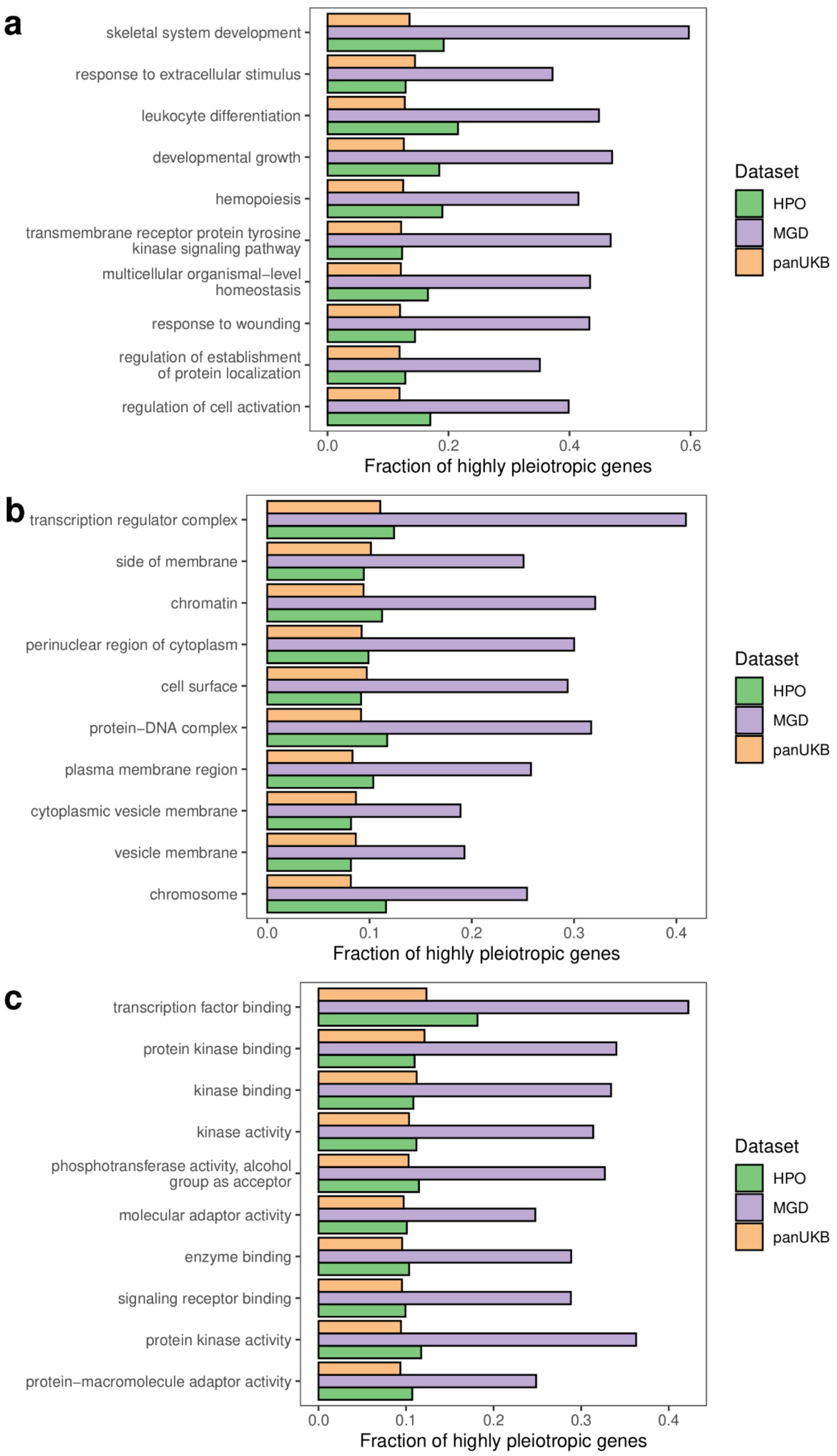


**Extended Data Figure 11. Shared processes, cellular localization and molecular functions of pleiotropic genes.** Shown are barplots of the fraction of highly pleiotropic genes among those annotated with a given biological process (a), cellular component (b), or molecular function (c) term from Gene Ontology. 10 terms with the highest proportions of pleiotropic genes are shown in each category.

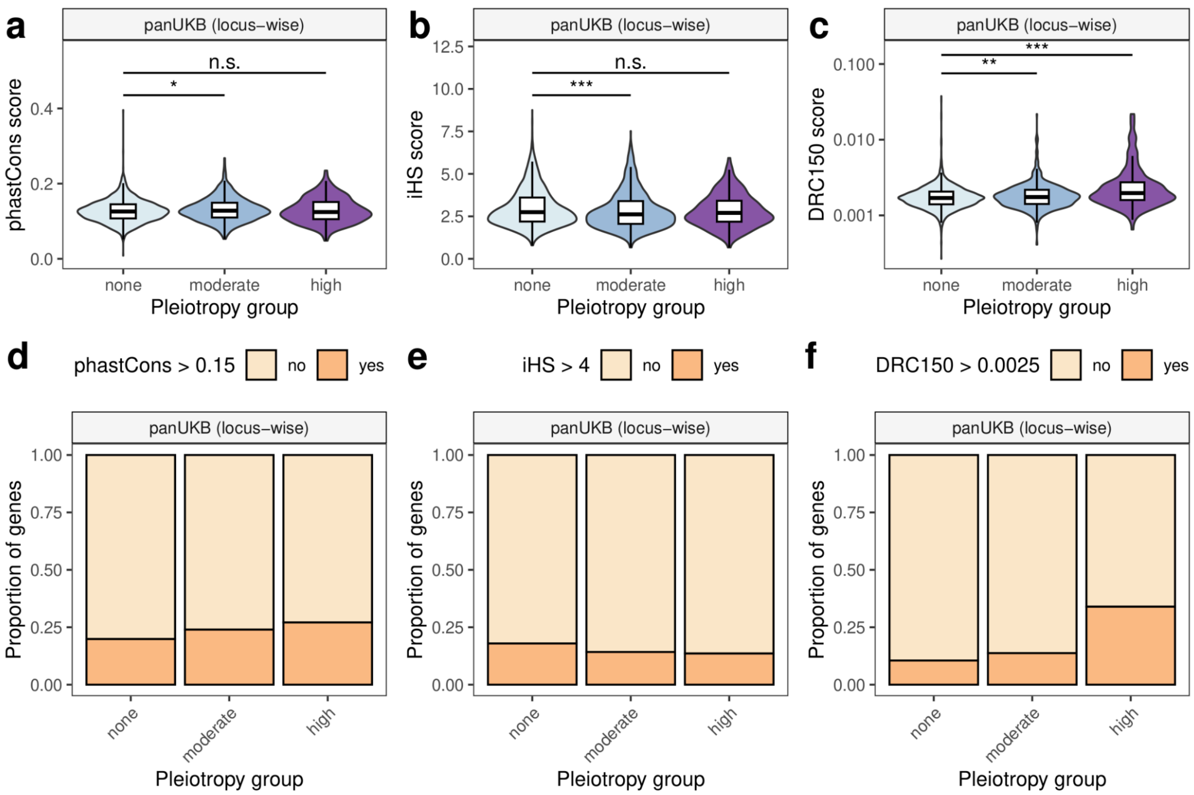


**Extended Data Figure 12. Locus-wise metrics of positive and negative selection in the pan-UKB GWAS data.** (a-c) Violin plots and box plots of average phastCons (a), maximum iHS (b) and DRC150 (c) values for each locus in the indicated pleiotropy groups. (d-f) Proportions of loci in the indicated groups that surpass a given cutoff for phastCons (d), iHS (e), or DRC150 (f).

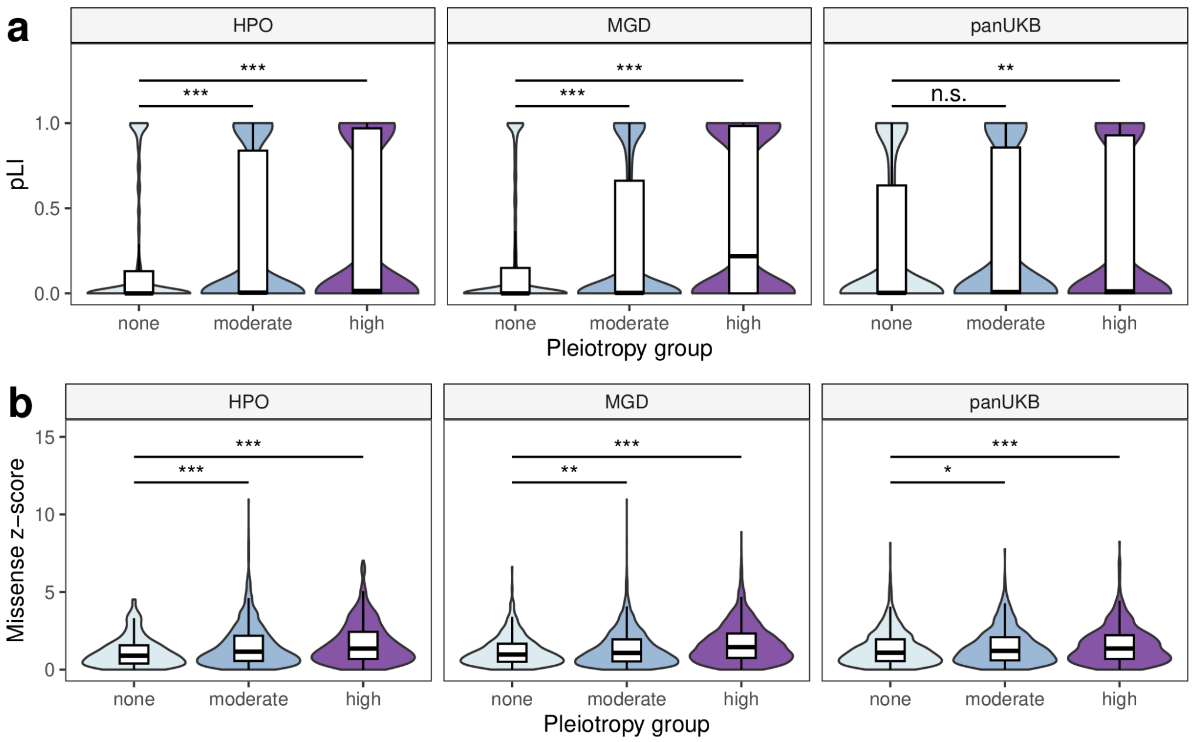


**Extended Data Figure 13. Highly pleiotropic genes have a high degree of constraint against both pLoF and missense variation.** Shown are violin plots and box plots of the probability of loss-of-function intolerance (pLI) and the Z-score of the observed-to-expected missense variant ratio (b) for indicated groups of genes. Values were taken from gnomAD v.2 data.

.

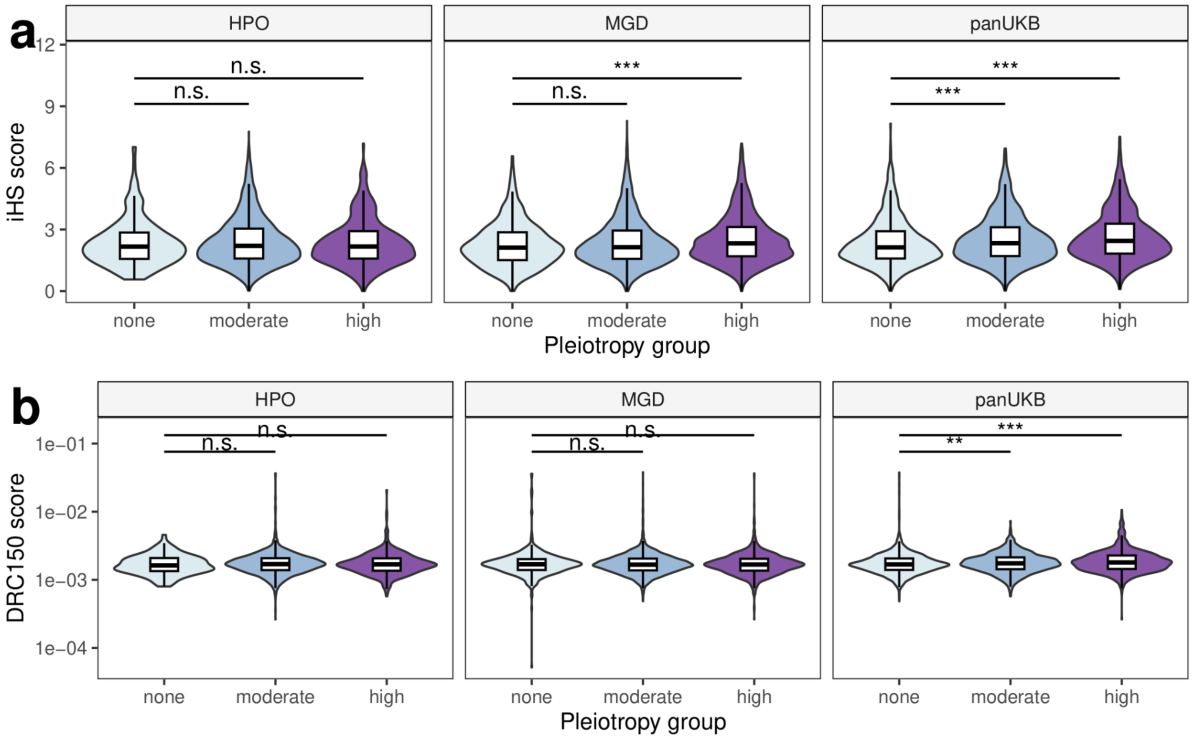


**Extended Data Figure 14. Highly pleiotropic loci show an enrichment with recent positive selection signals.** Shown are violin plots and box plots of the distribution of the gene-wise maximum iHS and DRC150 values for indicated gene groups.

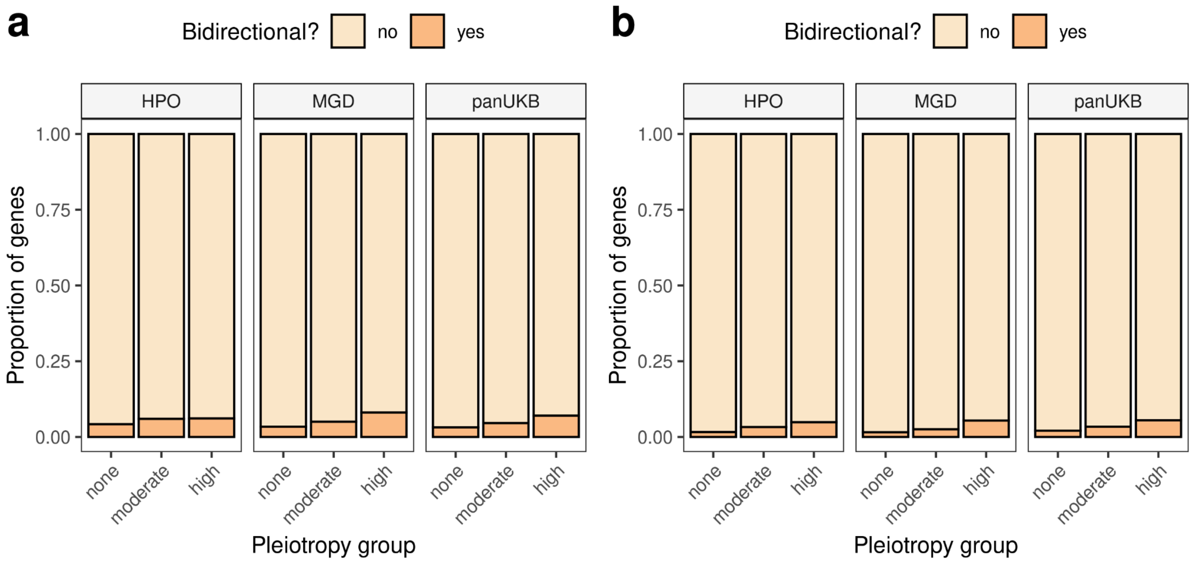


**Extended Data Figure 15. Highly pleiotropic genes bear combined signals of positive and negative selection.** Shown are bar plots representing the proportion of genes having a LOEUF value < 0.6 and either iHS > 4 (a) or DRC150 > 0.0025 (b).

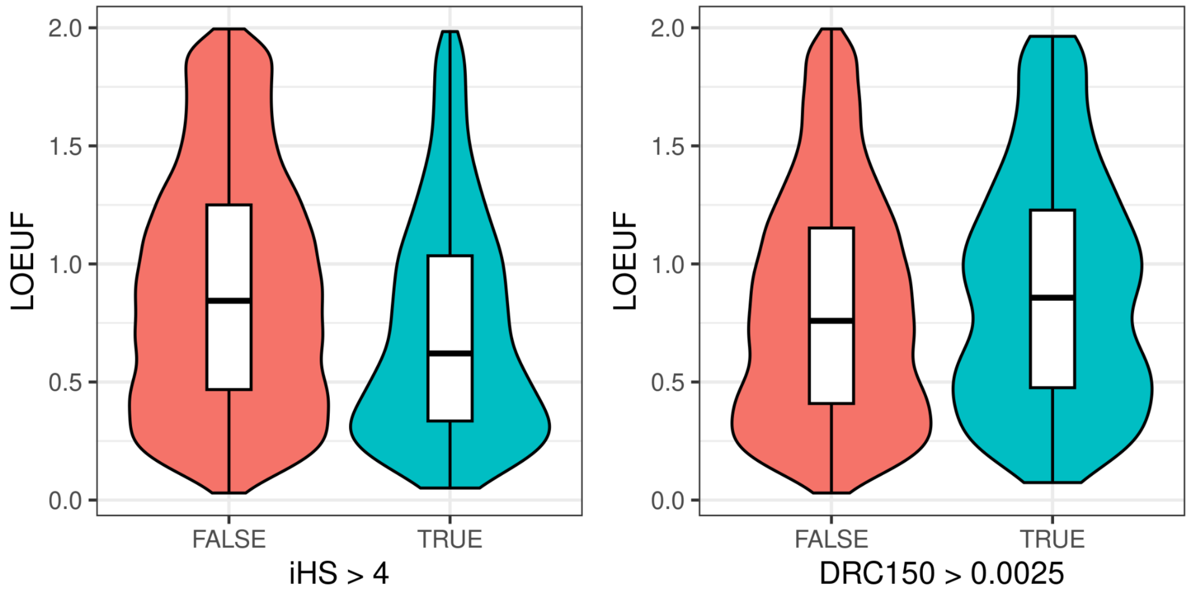


**Extended Data Figure 16. Statistics used for recent positive selection detection (iHS and DRC150) demonstrate different dependence on gene constraint.** Shown are violin plots and box plots of LOEUF values for genes that have lower (iHS <4, DRC150 < 0.0025) or higher (otherwise) values of recent positive selection statistics.

### Supplementary Note

In this Supplementary Note, we will discuss several important questions that were not addressed in the main text of the manuscript, but are of importance for understanding the methodology of the study.

#### 1. The choice of metric for assessing pleiotropy from phenotype ontology terms

Estimation of the *bona fide* (horizontal) pleiotropic effects is complicated by a variety of factors, one of which is correlations between traits. In the genome-wide association data, genetic correlation estimates enable grouping of traits with similar genetic basis to ensure that no highly correlated traits are counted as independent (see section 2 for a discussion of correlation-based clustering). For phenotype ontology term data gathered in monogenic trait setting, however, such estimates are not readily available.

Both Human Phenotype Ontology (HPO) and Mammalian Phenotype Ontology (MP) are hierarchically structured, with upper-level terms (e..g, "Abnormality of …") sometimes used as a proxy to measure pleiotropic effects (e.g., REFs). However, the distribution of gene-level term counts, while differing in the proportion of non-pleiotropic genes, have notable similarities in shape (Extended Data Figure 1). This poses the question regarding the validity of using upper-level PO terms as measures of horizontal pleiotropy. A similar question has been raised earlier in light of the Mouse Phenotype Consortium (MPC) data (REF Mammalian genome).

A possible way to remove redundancy in the data could be to use clustering of phenotype terms using data on associated genes. However, unlike complex trait clustering that can leverage genetic correlation estimates that are continuous in nature, clustering of phenotype ontology terms can be expected to suffer from a relatively low number of genes that are annotated with each specific term, thus making a distance metric used for clustering more discrete One might expect that, in such circumstance, unrelated terms could be spuriously grouped into a single cluster by virtue of having a single shared gene.

Nevertheless, we applied hierarchical clustering based on the Szymkiewicz–Simpson coefficient (overlap index) (a value of 1 - OI was used as a distance metric for clustering). The number of clusters was optimized using the silhouette score. Comparable number of clusters was identified for HPO and MP (5540 and 7138, respectively), with an average of 1.75 and 1.49 terms per cluster.

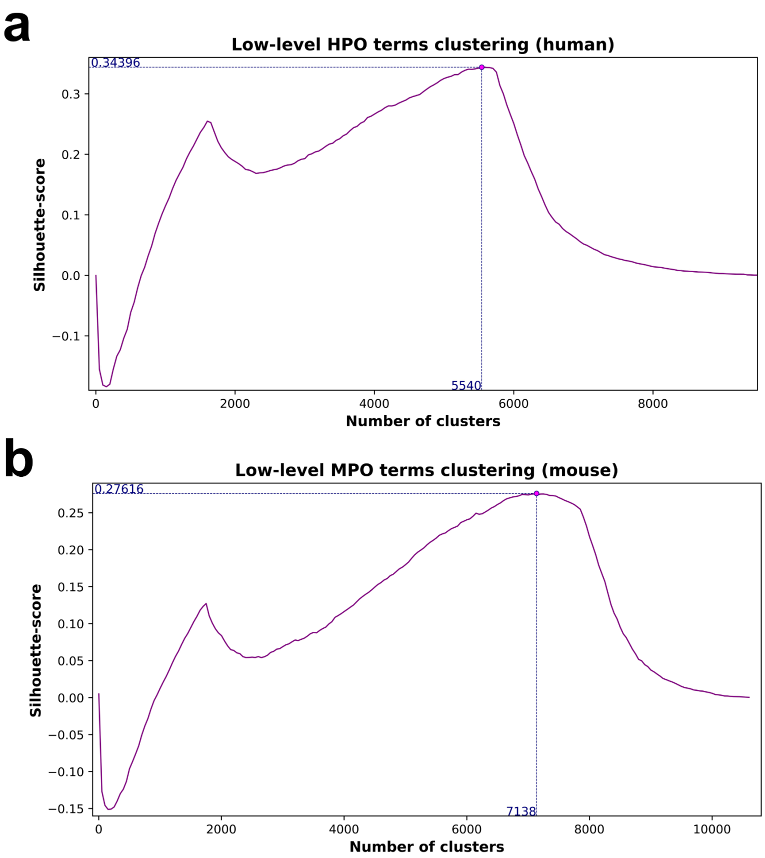


**Figure SN1.1.** Clustering of HPO and MP terms by similarity of the associated gene sets. Shown are line plots showing the silhouette score depending on the number of clusters for HPO (a) and MP (b) data.

We next used the obtained clustering of terms to re-calculate the gene-level degree of pleiotropy in both species. However, despite a shift in the distribution of the degree of pleiotropy when using clustered ontology terms compared to all ontology terms (Figure N1.2), the overall shape of the distribution was virtually unchanged in both human and mouse. The number of non-pleiotropic genes increased from 7 to 119 in humans, and from 909 to 1001 in mice. At the same time, this number was still smaller compared to the number of non-pleiotropic genes identified using upper-level MP terms as the measure of degree of pleiotropy (287 for humans and 1409 for mice). It is important to note that out of 119 genes with no pleiotropic effects according to the HPO term clusters, 76 (63.9%) remain non-pleiotropic according to upper-level MP term counts. For mice, this number amounts to 846 (84.5%).

Besides the number of non-pleiotropic genes, we also compared the median degree of pleiotropy per gene when using each of the approaches. Clustering of low-level ontology terms only slightly reduced the median pleiotropy compared to unclustered data in humans (from 40 to 33), and did not introduce any change for the murine data (in both cases, 10 terms or term clusters per gene were observed). Expectedly, the median degree of pleiotropy was much lower with upper-level terms as a measure (11 - for humans, and 5 - for mice).

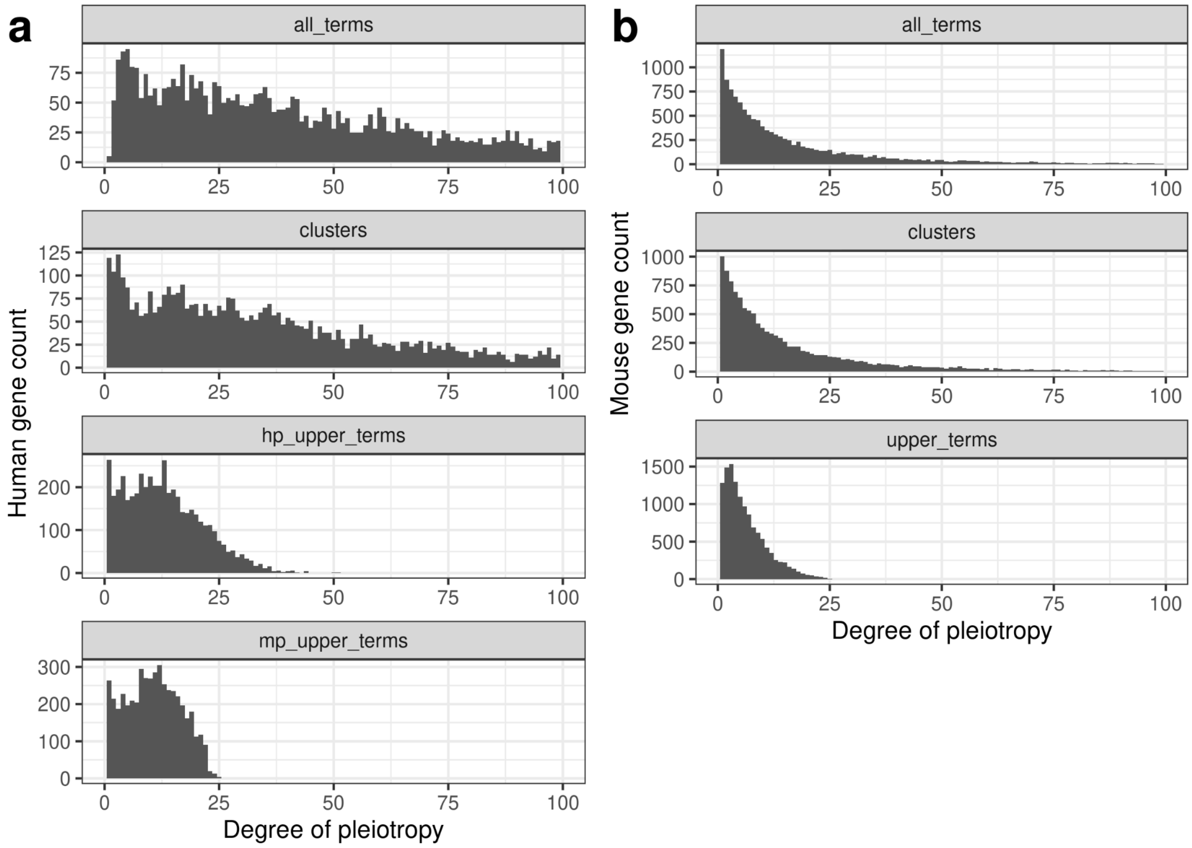


**Figure SN1.2.** The distributions of the per-gene degree of pleiotropy for human (left) and mouse (right) genes using either all ontology terms (top), or clusters of all ontology terms (middle), upper-level Mammalian Phenotype Ontology terms (bottom). For humans, an additional plot shows the number of upper-level HPO terms per gene before converting to MP vocabulary.

Besides the general similarity in the distribution shape, we have also evaluated the concordance between the number of upper-level HPO terms associated with a given gene and the number of matching upper-level MP terms after conversion. The results of the analysis showed that the values were expectedly highly correlated (Figure N1.3). More than 90% of genes (1013 out of 1109) that fell into the fourth quartile by their degree of pleiotropy using upper-level HPO terms were also in the fourth quartile of the distribution according to MP term count. The median degree of pleiotropy was also virtually the same (12 - with HPO terms, and 11 - with matching MP terms). Hence, we chose upper-level MP terms over upper-level HPO terms to enhance the comparison between the species.

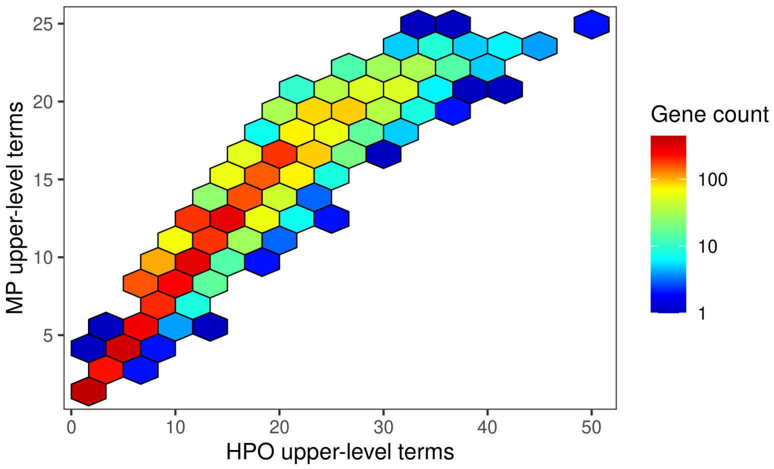


**Figure SN1.3.** Degree of pleiotropy estimated using upper-level HPO terms and mapped upper-level MP terms are strongly correlated. Shown is the hexagonal scatterplot of the number of terms associated with a given gene before and after converting HPO terms to MP terms. Color of the hexagon is proportional to the number of genes in the corresponding bin.

Taken together, these results suggest that clustering of ontology terms by similarity of the associated gene sets is less effective in reducing average degree of pleiotropy of a gene compared to grouping by upper-level terms. However, one has to bear in mind that this observation does not definitively prove that the upper-level terms are providing a better estimate of pleiotropy compared to clusters of lower-level terms.

**2. Do closest genes well represent the genotype-to-phenotype associations observed at GWAS loci?**

Another important issue pertinent to the analysis of gene-level pleiotropy from GWAS data is the identification of causal genes at the genome-wide significant loci. Three principal approaches can be utilized for solving this task: (i) picking the closest gene to each lead SNP identified after LD-based clumping of variants; (ii) using all genes that are located within a fixed size window from a lead SNP; and (iii) using statistical fine mapping or gene prioritization tools (reviewed in Uffelmann et al., 2021). The latter approach, however, is more complicated and relies on raw genotype information, making it hard to apply it to the summary data used in our work.

As expected, we found that much more genes appear as having at least one associated trait when using the second (locus-wide) approach compared to the closest gene approach (Figure SN2.1a). Thus, a total of 16,839 genes were identified using the locus-wide approach compared to only 6649 for the closest gene approach. Similarly, a median of 2 associations per gene were present in the former dataset, corresponding to 43.5% of non-pleiotropic genes (in closest gene-based data, a median of 1 association per gene was observed, and 56.4% of genes were thus non-pleiotropic).

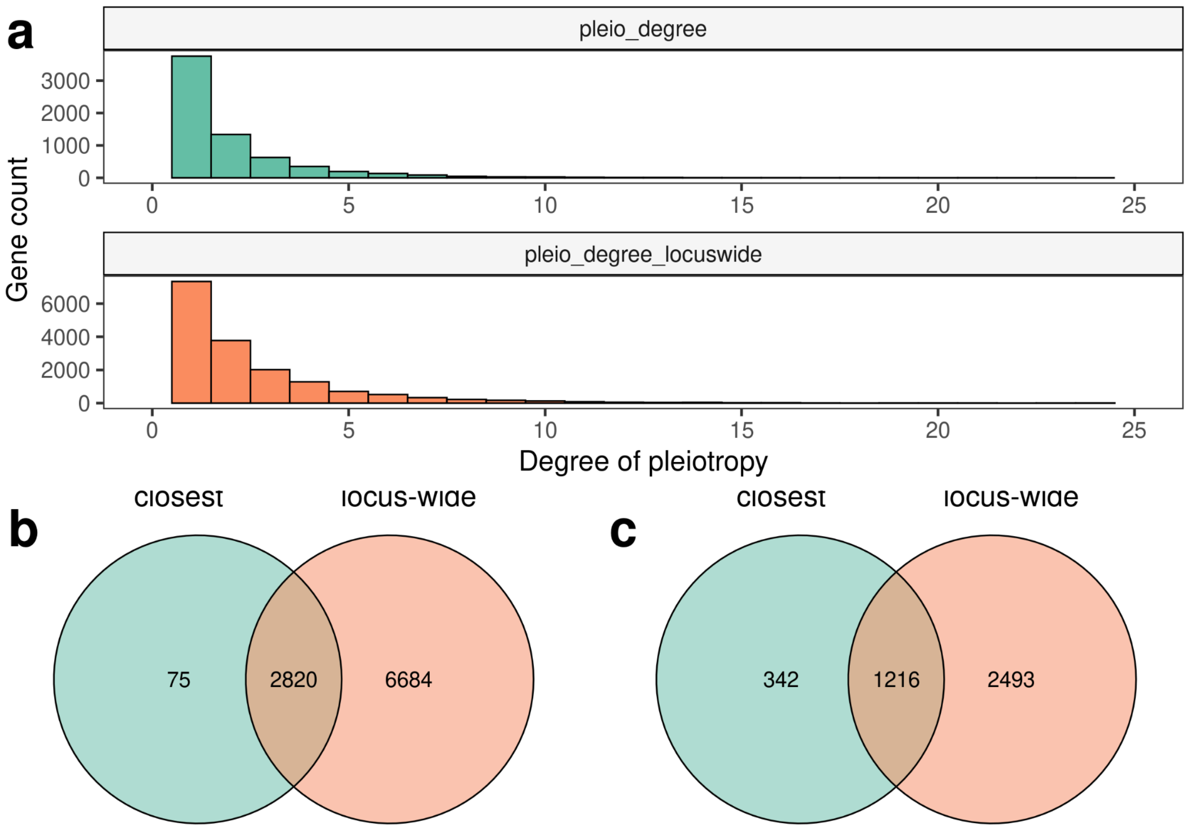


**Figure SN3.1.** (a) The distribution of the degree of gene-level pleiotropy calculated using closest gene-based and locus-wise (all genes within a 100,000 base pair window from lead SNP) GWAS data processing methods. (b, c). Venn diagrams showing the overlap between sets of pleiotropic (b) and highly pleiotropic (c) genes obtained using each method.

Expectedly, almost all of the genes that were identified as pleiotropic using the closest gene-based method were also pleiotropic in the locus-wise dataset (Figure SN2.1b). For highly pleiotropic genes, however, the overlap was lower, with as many as 342 genes designated as highly pleiotropic in the former dataset (Figure SN2.1c). This observation is likely driven by a lower cutoff for highly pleiotropic genes (3 trait clusters were required for closest gene-based data compared to 4 clusters for locus-wise data).

We next went on to evaluate whether the properties of pleiotropic genes identified using different data preprocessing strategies are similar, As shown in Figure SN2.2, the two strategies yielded similar results, with significant differences in expression pattern, GO term counts, and gene-level constraint (measured using LOEUF) observed in both datasets (Figure SN2.2a,c,d). Of note, highly pleiotropic gene set obtained using locus-wise method demonstrated a significant increase in the number of protein-protein interactions (Figure SN2,2b); however, the differences in LOEUF were, to the contrary, more pronounced when using closest genes (Figure SN2.2d). Notably, the difference in the expression profile for was more pronounced when all genes within a 100,000 kbp window from an index variant were used for analysis. For example, median expression of the genes at highly pleiotropic loci was 5.6-fold higher compared to non-pleiotropic genes (only a 2.7-fold difference was observed for closest genes). At the same time, highly pleiotropic genes closest to lead GWAS SNPs were expressed at 5 TPM in as many as 9 tissues (median) compared to only 2 tissues for all genes at a locus.

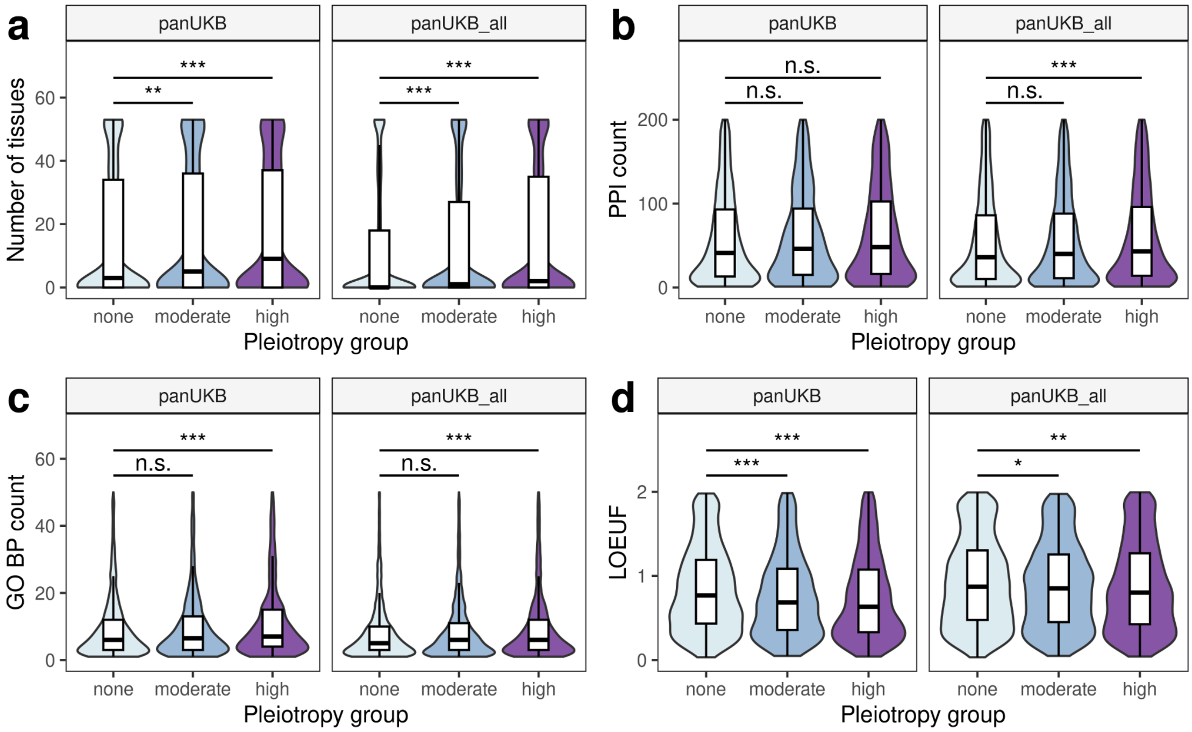


**Figure SN2.2.** Violin plots and box plots showing the distribution of gene properties for the given pleiotropy groups in GWAS data processed using closest gene picking ("panUKB") or taking all genes with a 100,000 bp window ("panUKB_all"). Shown are the number of tissues with expression at the level of > 5 TPM according to GTEx v.8 (a), the number of protein-protein interactions (b), the number GO biological process (BP) terms (c), and the loss-of-function observed-to-expected upper fraction (LOEUF) according to gnomAD v.2. (d). *** - $p < 0.001$, ** - $p < 0.01$, n.s. - no significant differences in the Wilcoxon-Mann-Whitney test with Benjamini-Hochberg FDR adjustment.

Taken together, the results presented above show that both locus-wise and closest gene-wisse analysis methods provide similar insights into the mechanistic basis of pleiotropic effects, and small differences observed when comparing the two datasets side-by-side can be attributed to the different cutoffs used for highly pleiotropic gene identification. The locus-wise approach, however, can be expected to yield a significant number of genes with false positive associations and spuriously pleiotropic genes that are located in the vicinity of several distinct causal genes. Hence, the closest gene-based approach was used in the main part of the analysis presented in the article.

#### 3. Pleiotropy analysis via clustering of complex traits compared to maximum independent set of traits

Previously, we have shown that clustering of traits by their phenotypic correlation is important to reduce spurious pleiotropy resulting from a large number of highly correlated traits in cohort datasets such as UK Biobank (Shukov et al., 2020). In particular, clustering allows the reduction of the proportion of the genome covered by pleiotropic loci from more than 60% (as reported by Watanabe et al., 2019) to ~5% (178.4 Mbp of the sequence).

In the pan-UK Biobank study (Karczewski et al., 2024), the authors have constructed the so-called maximum independent set of traits with low levels of correlation ($r^2 < 0.1$). In the data release used in our analysis, this set of traits comprised 150 traits of different types (). While such an independent set of traits should be perfectly suited to reduce the vertical pleiotropy in the data, one might expect that picking a single trait from a highly correlated cluster may significantly reduce the number of loci with genome-wide associations in the dataset. Hence, we compared the results of pleiotropy analysis using the maximum independent set of traits or the trait clustering used in the main part of our analysis.

First, we compared the distribution of the gene-level degree of pleiotropy in the two datasets. Expectedly, only 7092 gene-trait associations were detected using the maximum independent set, compared to 13,932 using the clustered dataset. The average degree of pleiotropy was slightly lower for the maximum independent trait set (1.74 compared to 2.10), and the proportion of pleiotropic genes was also slight decreased ((2895/6649, 43.5% compared to 1393/4072, 34.2%). The distribution of the degree of pleiotropy in the two data sources is shown on Figure SN3.1a. Besides, the degree of pleiotropy was highly correlated between the two datasets (Spearman's $\rho$ = 0.62 p < 0.001). 115 genes were uniquely pleiotropic in the maximum independent trait set (Figure SN3.1b), and only 83 genes were highly pleiotropic in the maximum independent set, with 592 highly pleiotropic genes overlapping between the two datasets (Figure SN3.1c).

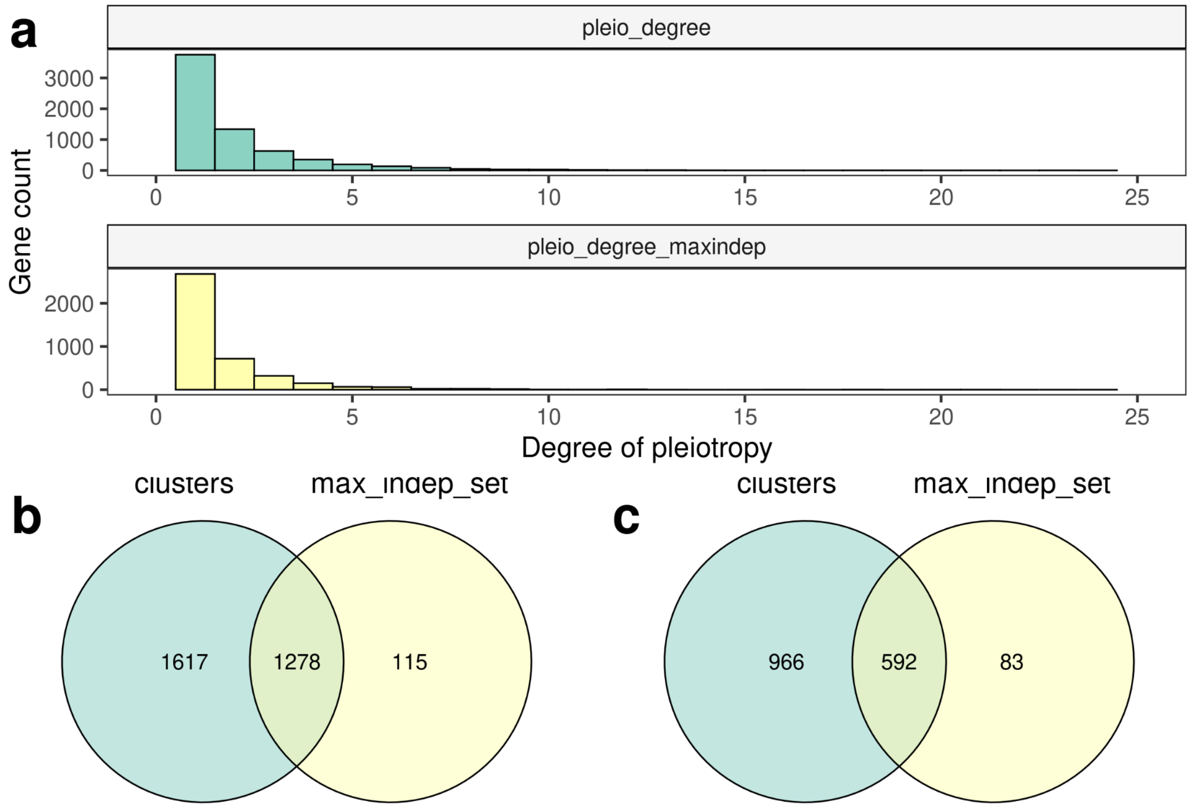


**Figure SN3.1.** (a) The distribution of the degree of gene-level pleiotropy calculated using clusters of correlated complex traits (top) or the maximum independent set of complex traits (bottom). (b, c). Venn diagrams showing the overlap between sets of pleiotropic (b) and highly pleiotropic (c) genes obtained using each method.

Given the subtle differences in the proportion of pleiotropic genes and the average degree of pleiotropy, we next moved on to test if the same differences in gene properties will be observed when using the maximum independent set of traits to measure pleiotropy in the complex trait domain. The analysis showed that, for all features that were evaluated, grouping of genes based on pleiotropy in the maximum independent trait set yielded the same results as the clustering-based approach (Figure SN3.2). Thus, significant differences remained between highly pleiotropic and non-pleiotropic genes in terms of the breadth of expression profile (number of tissues with gene expression at at least 5 TPM (Figure SN3.2a), number of biological process terms (Figure SN3.2c), and evolutionary constraint level (Figure SN3.2d)). For the number of protein-protein interactions, no significant differences were observed in either case (Figure SN3.2b). The only difference in results between the two analysis approaches was observed when comparing moderately pleiotropic genes to non-pleiotropic ones. For the expression profile metrics and gene constraint, pleiotropy estimation based on a maximum independent set did not allow to detect differences between these groups, unlike the approach based on trait clusters (Figure SN3.2).

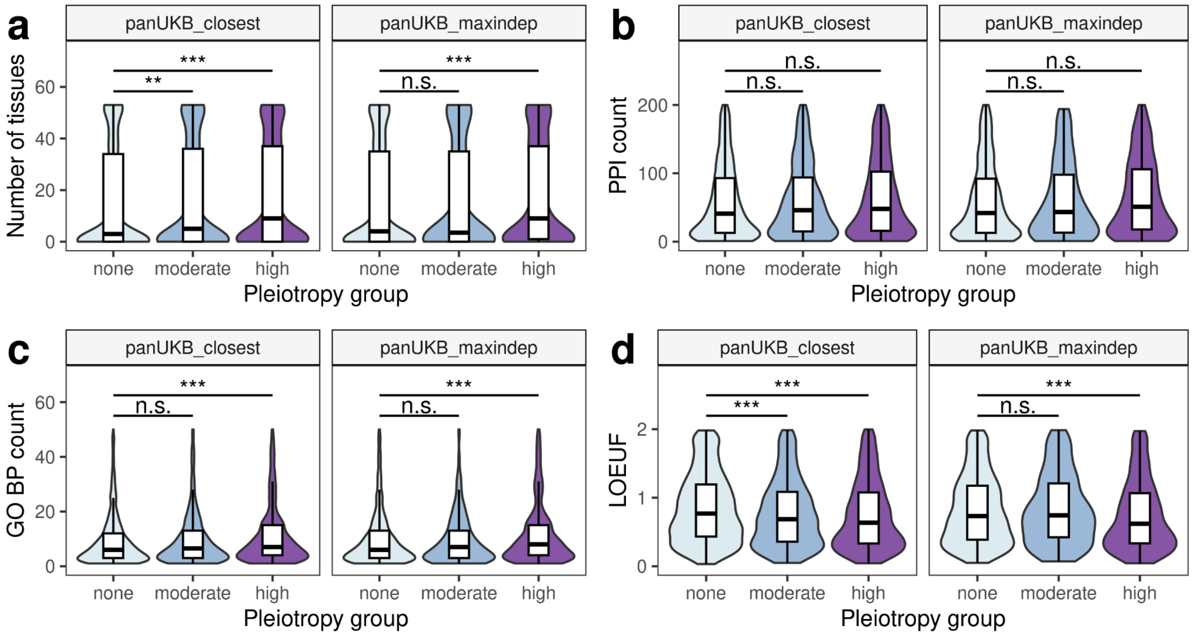


**Figure SN3.2.** Violin plots and box plots showing the distribution of gene properties for the given pleiotropy groups in indicated datasets. Shown are the number of tissues with expression at the level of > 5 TPM according to GTEx v.8 (a), the number of protein-protein interactions (b), the number GO biological process (BP) terms (c), and the loss-of-function observed-to-expected upper fraction (LOEUF) according to gnomAD v.2. (d). *** - $p < 0.001$, ** - $p < 0.01$, n.s. - no significant differences in the Wilcoxon-Mann-Whitney test with Benjamini-Hochberg FDR adjustment.

Taken together, the results presented in this section show that the maximum independent set of traits provided by the authors of the pan-UKB study does not provide any noticeable advantage over the approach based on trait clustering, and can even reduce the significance of certain trends. Hence, the method based on trait clusters was used as a standard one throughout the main part of the manuscript.
